# Theory on the accurate estimation of Michaelis-Menten enzyme kinetic parameters from steady state and progress curve datasets

**DOI:** 10.1101/2025.04.02.646753

**Authors:** Rajamanickam Murugan

## Abstract

We show that neither pure steady state nor pure progress curve analysis yield reliable estimate of enzyme K_M_ values since these methods are valid only at specific timescales and also use different type of datasets. All the currently proposed validity conditions of these methods assume *a priori* knowledge on K_M_. Hence, there is no way to check whether the obtained K_M_ from a given dataset is a reliable estimate or not. Here we propose an integrated approach in which the same time course dataset will be analysed both in the progress curve as well as steady state perspectives at different reaction timescales across replications and substrate concentrations. Our theory shows that there exists an optimum reaction time at which the error in the estimation of K_M_ using various progress curve and steady state methods show the least possible value so that the coefficient of variation of the median K_M_ values obtained across various methods attains a minimum. Using detailed stochastic simulations, we confirm that the K_M_ value obtained with minimum coefficient of variation across various methods is actually the reliable estimate that is close to the original K_M_ value. We further show that using multiple nonlinear regression methods, the type of inhibition viz. competitive, uncompetitive and mixed can be accurately classified apart from obtaining the accurate inhibitor constants and IC_50_ values from Dixon type average velocity steady state datasets.

## Introduction

Enzymes catalyse several crucial biochemical reactions especially at physiologically relevant timescales and environmental conditions [1-3]. The mechanistic view of single substrate enzyme catalysis can be well described by the Michaelis-Menten scheme (MMS) [4, 5]. In this scheme (**Fig. 1A**), the enzyme first binds the substrate to form an enzyme-substrate complex which reduces the activation energy barrier of substrate to product conversion. Subsequently, the substrate will be converted into the respective product and released. The MMS scheme is characterized by three different rate constants viz. diffusion-controlled bimolecular forward *k*_*1*_ (1/mol/lit/second), unimolecular reverse *k*_*-1*_ (1/second) and product formation rate constants *k*_*2*_ (1/second). These rate constants along with the initial enzyme level (e_0_) are generally summarized by two important enzyme kinetic parameters viz. *K*_*M*_ = (*k*_*-1*_ + *k*_*2*_) / *k*_*1*_ (mol/lit) and *v*_*max*_ = *k*_*2*_ e_0_ (mol/lit/s). Here k_2_ is also termed as k_cat_ and the ratio *k*_*E*_ = k_cat_ / K_M_ is defined as the catalytic efficiency of the enzyme. When *k*_*2*_ ≫ *k*_-*1*_, then k_E_ reaches the diffusion-controlled rate limit *k*_*1*_.

**FIGURE 1.**
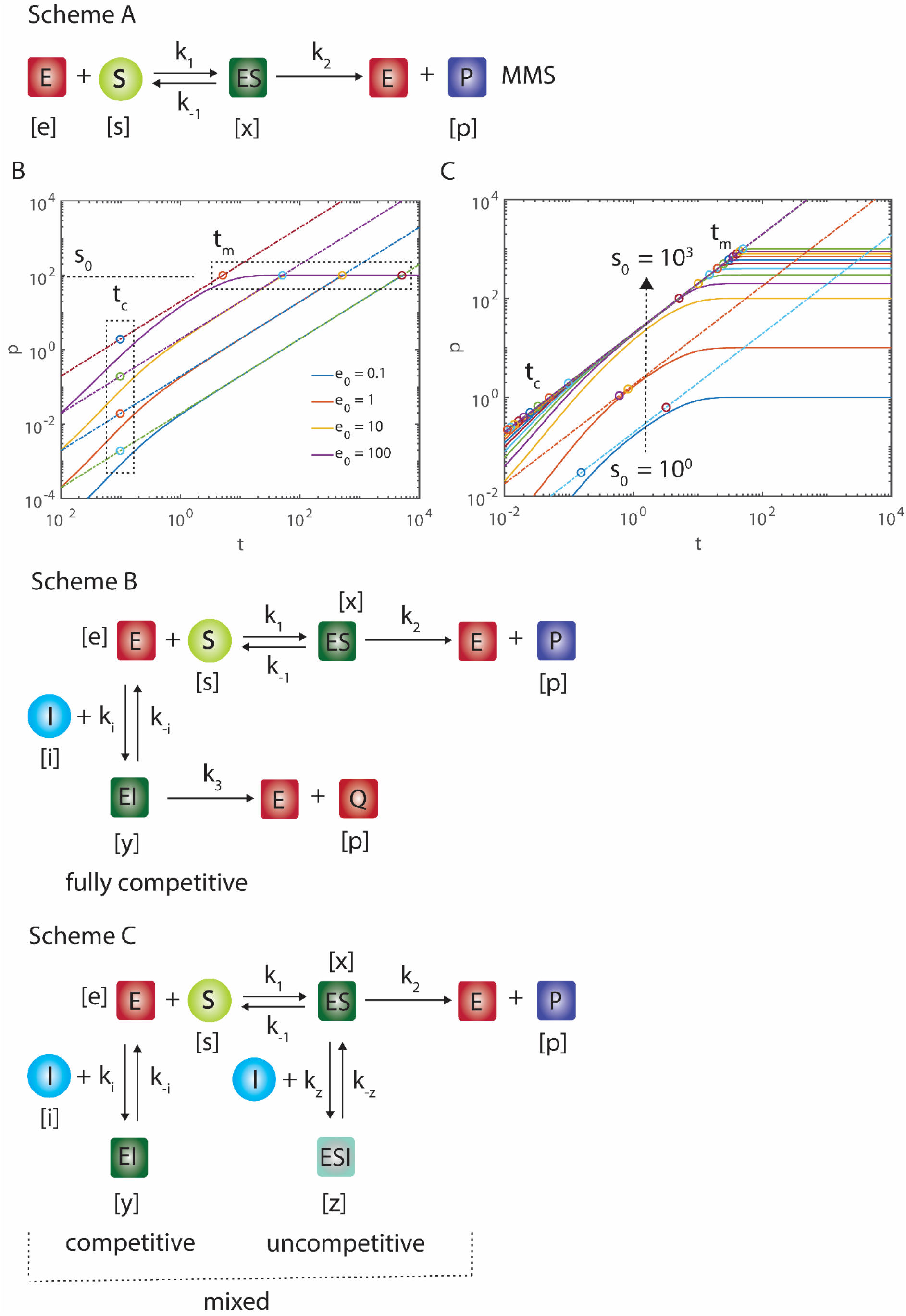
Dynamics of product formation (p, µM) in Michaelis-Menten scheme of single substrate enzyme catalyzed reactions. Here *t*_*c*_ ≅ 1⁄*k*_1_(*K*_*M*_ + *s*_0_), *t*_*m*_ ≅ ((*K*_*M*_ + *s*_0_)⁄*k*_*2*_*e*_0_) are the approximate pre-steady state and reaction-end timescales (t, seconds) and numerical integration settings from **Eqs. 1** are k_1_ = 0.1 µM^-1^s^-1^, k_-1_ = 0.01 s^-1^ and k_2_ = 0.2 s^-1^, with K_M_ = 2.1 µM. **A**. *s*_0_ = 100 µM and e_0_ was iterated in (0.1, 1, 10, 100) µM. **B**. *e*_0_ = 100 µM and s_0_ was iterated in (1, 10, 100-1000, with increment of 100) µM along the arrow. Scheme B describes the fully competitive inhibition and Scheme C describes various partial inhibition schemes viz. competitive (k_z_ = 0), uncompetitive (k_i_ = 0) and mixed types. In fully competitive inhibition both substrate and inhibitor will be converted into their respective products by the same enzyme. Whereas, in case of partial competitive inhibition the enzyme-inhibitor will be a dead-end complex.

Exact closed form analytical solution to the nonlinear set of ODEs corresponding to MMS scheme is not known though the integral solution can be expanded in terms of ordinary [6] and singular perturbation series [7] or over slow manifolds [8, 9]. Singular perturbation expansions always yield inner and outer solutions corresponding to the pre-steady state and post-steady state regimes which should be combined via proper matching at the temporal boundary layer [7, 10-15] to obtain the complete solution. Whereas, ordinary perturbation series can give the complete solution that is valid for both pre- and post-steady state regimes [6]. Obtaining (*K*_*M*_, *k*_*cat*_, *v*_*max*_) from the steady state and progress curve experimental datasets is essential to characterize an enzyme. Since complete closed form analytical solution to MMS is unknown, one needs to rely on the widely used standard quasi steady state approximation (sQSSA) of the reaction velocity *v* = d*p*/d*t* as *v* = *v*_*max*_ *s* / (*K*_*M*_ + *s*) that is valid only when k_2_ ≪ (k_1_ s_0_) [16]. Here p represents the product concentration. When e_0_ ≪ s_0_, then one can use the stationary reactant assumption to further simplify as v ≅ v_0_ = v_max_ s_0_ / (K_M_ + s_0_) which connects K_M_, v_max_ and initial substrate concentration s_0_. Nonlinear least square fitting of the data on (v, s_0_) over original hyperbolic function or linearization techniques such as Lineweaver-Burk on (1/v, 1/s_0_) and Eadie-Hofstee on (v, v/s_0_) [17] can be used to obtain the estimates of kinetic parameters [18, 19]. Several statistical analysis procedures were also proposed [20, 21] to compute the error on these parameter estimates.

Enzymes of pathogenic organisms are important targets for drug molecules to control various diseases. Accurate estimation of various MMS parameters especially K_M_ along with K_I_ of a competitive inhibitor is essential to compute the IC50 values of drug molecules against a given target enzyme [22-25]. In steady state experiments, a fixed amount of enzyme will be incubated with a series of different substrate concentrations (s_0_) for fixed amount of time (t_r_) at constant temperature and pH conditions from which the *average product formation velocity* 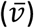 will be calculated by the ratio = total product formed during incubation time / total incubation time (t_r_). Generally, t_r_ will be fixed approximately at 1 % depletion of s_0_ however without any theoretical basis. Under sQSSA conditions with e_0_ ≪ s_0_, it will be approximately assumed that the system evolves with constant reaction velocity *v*_0_ = v_max_ s_0_ / (K_M_ + s_0_) after a short pre-steady state timescale *t*_*c*_ ≪ *t*_*r*_ so that the total product formed over time t_r_ will be approximately equal to (p = v_0_ t_r_) and therefore one sets the average reaction velocity 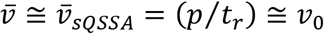. This equation is being used widely across enormous volumes of literature to estimate K_M_ and v_max_ from the dataset on 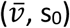 obtained from steady state experiments. To estimate the inhibition constants K_I_, this average velocity experiments will be conducted at different inhibitor concentrations i_0_, and the data on (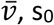, s_0_, i_0_) will be used to classify the type of inhibition (i.e. competitive, uncompetitive or mixed type) and estimate v_max_, K_M_ and K_I_ using Dixon plots [26] or sequential or simultaneous nonlinear least square fitting procedures [27, 28]. Using these parameters IC_50_ value of the inhibitory drug will be computed using Cheng-Prusoff equation [23]. Here 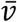 is the average velocity = product formed in a given total reaction time / total reaction time.

Detailed studies proposed the essential and sufficient conditions for the validity of the approximation 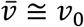 viz. (a) k_2_ ≪ (k_1_ s_0_) [16] (b) e_0_ ≪ s_0_ [29, 30] (c) (e_0_ / (s_0_ + K_M_)) ≪ 1 [29, 30]. However, except (b), conditions (a) and (c) strictly require *a priori* knowledge on k_1_, k_2_ and K_M_. Otherwise, one needs to set very high values of s_0_ along with condition (b) so that all the conditions (a-c) are satisfied. Although the inequality (b) is possible under *in vitro* laboratory conditions, there are several situations such as single molecule enzyme kinetics and other *in vivo* conditions where one cannot manipulate the ratio of substrate to enzyme concentrations much. Setting high s_0_ along with infinitesimal e_0_ ≪ s_0_ would eventually drives 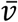 towards infinitesimal values below the background noise level which results in noisy experimental data on (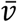, s_0_) that leads to enormous amount of error in the estimation of K_M_. Several recent studies [32-35] reported this issue and also put forth alternate methods such as total QSSA (tQSSA) to obtain accurate K_M_ values from such experimental datasets.

The source datasets for the estimation of K_M_ will be either time dependent progress curve type [36] or steady state average velocity type. Here progress curve data on (p, s, t) contains information on both pre-as well as post steady state dynamics that in turn depends on the timescale of experimental data collection. Whereas, steady state experimental setups generally capture the details of only the post-steady state regime. Progress curve methods can yield accurate estimates of K_M_ only when the data collection done at those timescale regimes with maximum change in the curvature of the kinetic trajectory [37]. Further, the role of total reaction time t_r_ is not considered anywhere in the sQSSA calculations i.e., 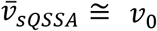though the final results strongly depend on t_r_. In this context, integrated form of sQSSA (isQSSA) is more superior than the general sQSSA since it also considers t_r_.

Since the conditions of validity of various methods like sQSSA, isQSSA and progress curve analysis are different from each other, K_M_ values of the same enzyme obtained from each of these methods under identical conditions using the same source dataset will not be consistent. Further, absence of *a priori* knowledge on K_M_ and other kinetic parameters will eventually introduces uncertainty in setting up the appropriate experimental conditions to minimize the error in the estimation of these parameters using sQSSA methods with stationary reactant assumption or progress curve methods. Apart from the sequential and simultaneous nonlinear fitting, analysis of Dixon type inhibition datasets on (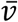, s_0_, i_0_) warrants multiple nonlinear regression methods (MNLR) to obtain reliable estimates of IC_50_ of drug molecules. In this article, we develop an integrated theoretical and computational approach which can produce consistent estimate of v_max_, K_M_, K_I_ and IC50 upon considering all these uncertainty aspects.

## Theoretical Methods

### Single substrate Michaelis-Menten kinetics

The nonlinear coupled differential rate equations corresponding to MM **Scheme A** shown in **Fig. 1** can be written as follows [16].

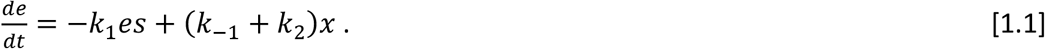

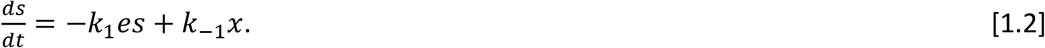

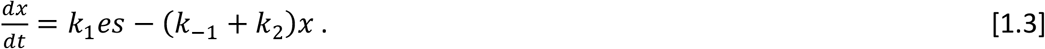

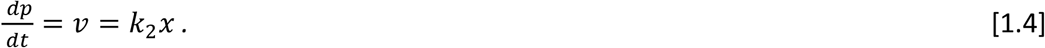

Here k_1_ (1/mol/lit/second), k_-1_ (1/second) and k_2_ (1/second) are the rate constants corresponding to bimolecular enzyme-substrate complex formation, dissociation and product formation respectively. Further, (*e, s, x, p*) (mol/lit) are the concentrations of enzyme, substrate, enzyme-substrate complex, product respectively and v (mol/lit/second) is the reaction velocity. The initial conditions at *t* → 0 are (*e, s, x, v, p*) = (*e*_0_, *s*_0_, 0,0,0) and the reaction trajectory ends at (*e, s, x, v, p*) = (*e*_0_, 0,0,0, *s*_0_) as *t* → ∞. The dynamical variables (*e, s, x, p*) obey the mass conservation laws viz. *s* = (*s*_0_ − *x* − *p*), *e* = (*e*_0_ − *x*) where one can also replace *x* = *v*⁄*k*_*2*_. Here the required enzyme kinetic parameters are defined as *K*_*M*_ = (*k*_−1_ + *k*_*2*_)⁄*k*_1_ and *v*_*max*_ = *k*_*2*_*e*_0_. From the definition of K_M_, one can conclude that accurate estimation of K_M_ requires detailed data points representing both pre- and post-steady state dynamics. To simplify the system of **Eqs. 1**, we first introduce the following set of scaling transformations [16].

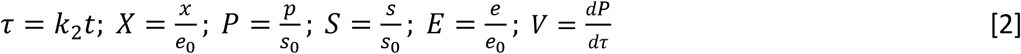

In **Eqs. 2**, the variables (e, s, x, p) are normalized so that 0 ≤ (*E, S, X, P*) ≤ 1 with the mass conservations laws *S* = 1 − *εX* + *P, E* + *X* = 1. We define *V* = *εX* so that *V* + *P* + *S* = 1 Where 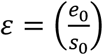. **Eqs. 1** can be rescaled in multiple ways at different variable spaces by the following set of dimensionless nonlinear ODEs [16].

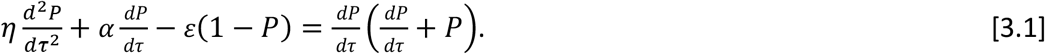

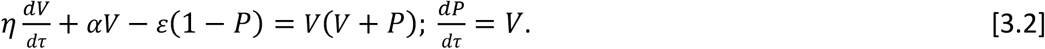

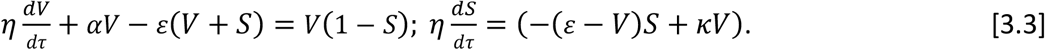

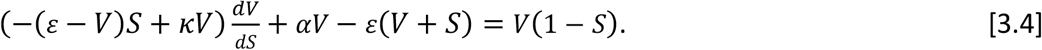

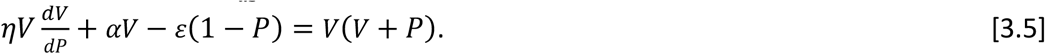

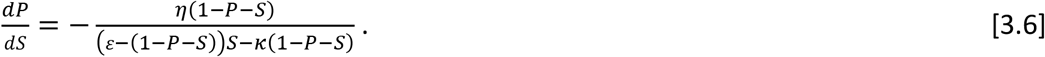

Here **Eq. 3.1** describes the MMS dynamics in (P, τ) space, **Eqs. 3.2** in (V, P, τ), **Eqs. 3.3** (V, S, τ), **Eq. 3.4** in (V, S), **Eq. 3.5** in (V, P) and **Eq. 3.6** in (P, S) spaces. Various dimensionless parameters in **Eqs. 3** are defined as follows.

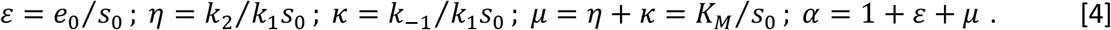

When *η* → 0, then **Eq. 3.4** results in sQSSA with the velocity *v*_*sQSSA*_ in the original (v, s) space. Similarly, **Eq. 3.5** results in tQSSA with the velocity *v*_*tQSSA*_ in the (v, p) space which can be written in terms of the original dynamical variables as follows.

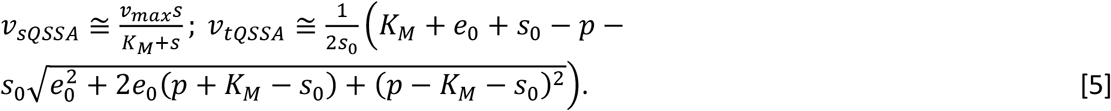

We denote **Eqs. 5** as *η*-approximations since they are valid only when *η* → 0. The main flaw of these sQSSA as well as tQSSA methods is that the left-hand sides of **Eqs. 5** are still time dependent quantities along with time dependent s and p on the right-hand sides. That is to say, *v*_*sQSSA*_ and *v*_*tQSSA*_ are tangents of progress curve in (p, t) space at a given time point and not the time averaged velocities as used in most of the steady state experiments. To derive equations which are useful to fit steady state experimental data, one needs to further set *ε* → 0 in **Eqs. 5**. This leads to the stationary reactant assumption *s* ≅ *s*_0_ − *p* ≅ *s*_0_ that results in the constant velocity as 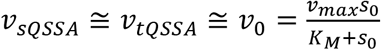 since *lim*_*ε*→0_*v*_*tQSSA*_ ≅ *v*_*sQSSA*._ In this scenario, one can conduct the average steady state velocity experiments only within the reaction timescales in which these conditions are valid.

Upon setting *ε* → 0 in **Eq. 3.6**, one finds that 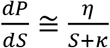 in the pre-steady state regime of (P, S) space. Solving this ODE for the initial condition P = 0 for S = 1, we obtain 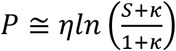. Since *S* ≅ 1 when (*η, ε*) → (0,0), one finds that *P* ≅ 0 and *V* ≅ 1 − *S* in the pre-steady state regime (*p* ≅ 0 and *v* ≅ *k*_*2*_(*s*_0_ − *s*) in terms of original variables). Since *s* ≅ *s*_0_ when *s*_0_ ≫ *s*_0_, one finds that *v* ≅ 0 for infinitesimal values of *k*_2_ near the steady state regime. The appropriate working reaction timescales for the average steady state velocity experiments can be derived as follows. Using the transformation rule *P* = *εη*^*2*^*U*, **Eq. 3.1** can be rewritten as the following ordinary perturbation equation with the scaling parameter *ϕ* = *ηε* as follows.

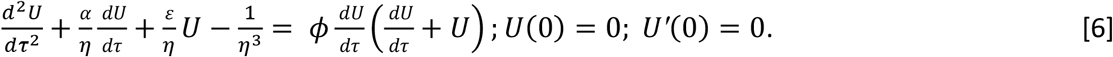

When *ε* → 0, then one arrives at the following approximations from **Eqs. 6** using the definition of dimensionless velocity *V* = *εη*^*2*^*F*.

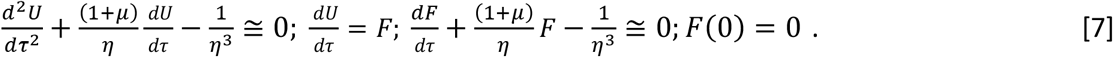

We denote **Eqs. 7** as *ε*-approximations. Solving **Eqs. 7** for (U, F) in terms of τ and then reverting back to (P, V) and subsequently transforming back in to the original variables (p, v, t), one arrives at the following ε-approximations.

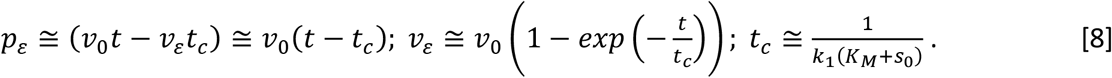

Here 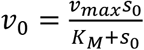 is the sQSSA with stationary reactant assumption i.e., *s* ≅ *s*_0_. When *t* ≫ *t*_*c*_, then one finds that *p*_*ε*_ ≅ *v*_0_(*t* − *tc*). From **Eqs. 8** one can also obtain *s*_*ε*_ ≅ *s*_0_ − *v*_0_*t* −*v*_*ε*_ (1⁄*k*_*2*_ − *t*_*c*_) which follows from the mass conservation law *s*_*ε*_ ≅ *s*_0_ − *p*_*ε*_ − *v*_*ε*_⁄*k*_*2*_. The subscript ε in *p*_*ε*_ and *s*_*ε*_ corresponds to the ε-approximations. The pre-steady state timescale in **Eq. 8** is *t*_*c*_. When *t* ≫ *t*_*c*_ then one finds that *s*_*ε*_ ≅ *s*_0_ − *v*_0_(1⁄*k*_*2*_ − *t*_*c*_) − *v*_0_*t*. When *t* =*t*_*c*_, then the steady state concentration of substrate can be approximated as *s*_*εc*_ ≅ *s*_0_(1 − *e*_0_⁄(*K*_*M*_ + *s*_0_)). Clearly, the stationary reactant assumption [38] *s*_*εc*_ ≅ *s*_0_ will be true only when (*e*_0_⁄(*K*_*M*_ + *s*_0_)) ≪ 1 which will be eventually true when *ε* → 0. When the reaction time t_r_ is such that *t*_*r*_ *< t*_*c*_, then *p*_*ε*_ ≅ 0 in the pre-steady state regime (infinitesimal quantity especially for *ε* → 0). When *t*_*r*_ ≫ *t*_*c*_, then one finds the linear product evolution regime with reaction time t_r_ as *p*_*ε*_ ≅ *v*_0_(*t*_*r*_ − *t*_*c*_) along with the constant reaction velocity *v*_*ε*_ ≅ *v*_0_. However, this linear dependence of *p*_*ε*_ on reaction timescale will break down towards the end of reaction since the substrate concentration starts to decline towards zero beyond this timescale *t*_*m*_. This approximately occurs when 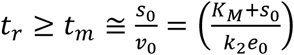. Since the product concentration evolves linearly with time as *v*_0_*t* in the post-steady state regime, we find that 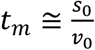. These timescale behaviors are demonstrated using numerically computed integral trajectories of **Eqs. 1** in **Fig. 1B** and **1C**. Clearly, one finds that 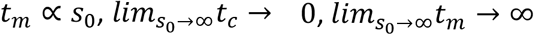 and 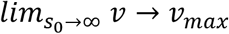 Further, one can define the timescale separation ratio *χ* as follows.

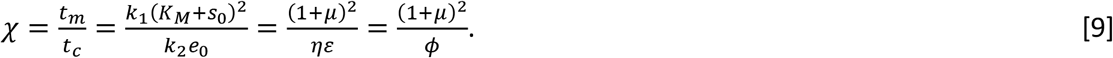

Here *χ* is inversely proportional to the perturbation parameter *ϕϕ*. The successfulness of sQSSA methods strongly depends on extent of timescale separation that is decided by the condition *χ* ≫ 1 [16, 30] which also means that *ϕϕ* = *ηε* ≪ (1 + *µ*)^*2*^. The linear dependence of product formation on time will be valid only when the reaction time falls in the range *t*_*c*_ ≪ *t*_*r*_ *< t*_*m*_ where one can conduct average reaction velocity experiments using 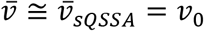. However, fitting of steady state data on 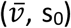with *v*_0_ will be biased towards v_max_ over K_M_ since there is not enough datapoints from the pre-steady state regime. When *t*_*r*_ *< t*_*m*_, then one can also use **Eqs. 8** to conduct average velocity experiments in the following form.

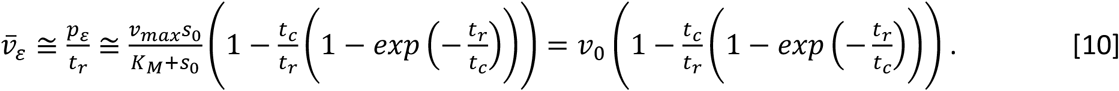

Here *p*_*ε*_ is the total product formed at the end of reaction time t_r_ as in **Eq. 8** and 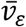 is the average velocity obtained from the steady state experiments. Unlike 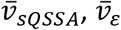 also considers the nonlinearity of the pre-steady state regime. The conditions for the validity of **Eq. 10** are

*ε* → 0 and *t*_*r*_ *< t*_*m*_. When *t*_*m*_ > *t*_*r*_ ≫ *t*_*c*_, then 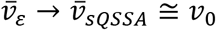. Here, 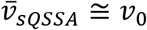 directly connects the average reaction velocity with the total substrate concentration s_0_. Clearly, the conditions for the validity of the widely used 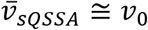 are viz. (*η, ε*) → (0,0) and *t*_*c*_ ≪ *t*_*r*_ *< t*_*m*_. Nonlinear least square fitting of the data on 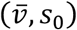, for a given reaction time t_r_ with **Eq. 10** can directly yield the parameters (K_M_, v_max_ and k_1_). However, such fitting procedures favour accurate estimation of v_max_ over (K_M_, k_1_) since they mainly rely on the datapoints from the pre-steady state timescale regime which is actually a transient one compared to the post-steady state timescale. To overcome this issue, we will fix k_1_ in the definition of 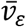 with the median k_1_ obtained from the progress curve methods in the later sections. To minimize the error in the estimation of K_M_ one needs to set *t*_*r*_ such that it is much lesser than the maximum reaction time t_m_ corresponding to the lowest s_0_ value used in the entire experimental setup. When *η* → 0, then one finds as in **Eqs. 5** that 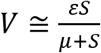. When *ε* → 0, then one finds that 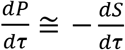 in the post-steady state regime and therefore one obtains 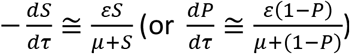 Upon solving this nonlinear first order ODE with the initial condition S = 1 at τ = 0, and then reverting back to the original dynamical variables, one finds the following implicit solution.

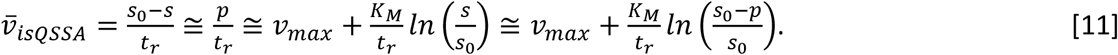

**Eqs. 10** is another form of integrated rate equation [35] that directly connects the average reaction velocity 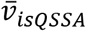 with the enzyme kinetic parameters and reaction time *t*_*r*_. Data on 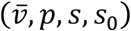, at a given reaction time *t*_*r*_ can be directly used to perform linear least square fitting with **Eq. 11** i.e., 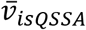 versus 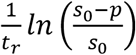 (type I) or 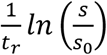 (type II) to obtain K_M_ and v_max_. In the later sections, we will show that type II form gives more accurate estimate of K_M_ than type I. The conditions for the validity of **Eq. 11** are (*η, ε*) → (0,0) and *t*_*c*_ *< t*_*r*_ *< t*_*m*_. One can also integrate 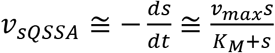 given in **Eqs. 5** under the conditions *ε* → 0 for the initial condition s = s_0_ at t = 0 and derive the following expression for the average reaction velocity.

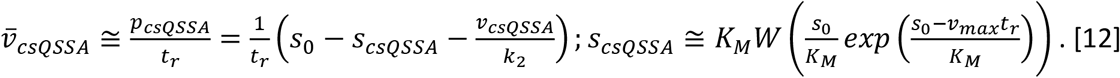

In this equation, we have used the mass conservation law *p* = (*s*_0_ − *s* − *v*⁄*k*_*2*_) and 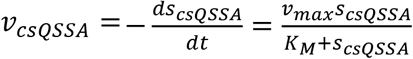. When 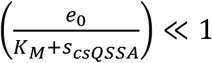, then one finds from **Eqs. 12** that 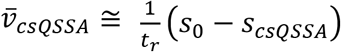 where *p*_*csQSSA*_ ≅ (*S*_0_ − *S*_*csQSSA*_) which is an alternate form on **Eq. 11**. In **Eqs.12**, W(x) is the Lambert W function [39, 40] which is the solution of y exp(y) = x for y. The conditions of validity of **Eqs. 12** is similar to **Eqs. 11**. We use **Eq. 11** for comparison purposes since **Eq. 11** is simple and computationally cheaper than **Eq. 12**. K_M_ and v_max_ can be obtained from the steady state datasets using any one of the expressions 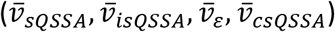 under appropriate validity conditions and timescales. However, except the condition on *ε*, all the others conditions (on *η*, t_c_, t_m_) require *a priori* knowledge on k_1_, k_2_ and K_M_ which are generally not available for unknown enzyme systems. In this scenario, one can perform direct non-linear least square fitting of the time course kinetic data on (p, t) or (s, t) with the system of nonlinear ODEs given in **Eqs. 1** to obtain k_1_, k_-1_ and k_2_. However, accuracy of such fitting procedure strongly depends on the detailed data points on both pre- and post-steady state regimes that warrants progress curve data collection on sub-millisecond timescales apart from the steady state timepoint.

### Michaelis-Menten kinetics with competitive inhibition

When (*ε*, *η*, *η, ε*) → (0,0,0,0) and 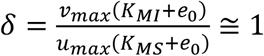, then earlier detailed theoretical studies revealed [22] the following approximations for the average velocities of the fully competitive inhibition **Scheme B** described in **Fig. 1**.

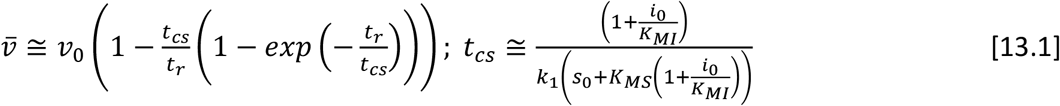

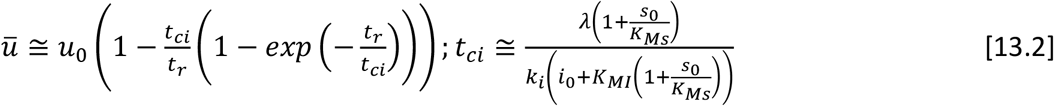

In fully competitive inhibition, same enzyme will catalyse the conversion of two different substrates (s, i) in to their respective products (p, q) with reaction velocities (v = dp/dt, u = dq/dt). In these equations, various parameters as defined as follows.

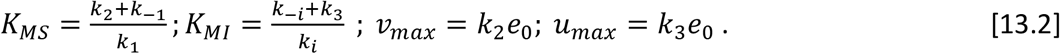

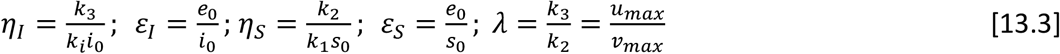

When the total reaction time is such that *t*_*r*_ ≫ *t*_*cs*_ and *t*_*r*_ ≫ *t*_*ci*_ where (*t*_*cs*_, *t*_*ci*_) are the pre-steady state timescales, then **Eqs. 13.1-2** reduce to the following well-known steady state velocity equations.

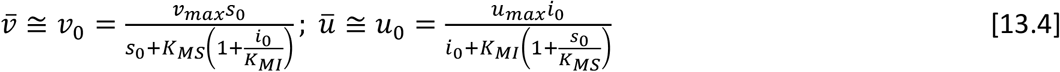

When δ ≪ 1 or δ ≫ 1, the validity of **Eqs. 13.4** will break down since the fully competitive inhibition system can exhibit multiple steady states [22] under such scenarios. Similarly for pure partial competitive inhibition given **Scheme C** of **Fig. 1**, one can derive the following average velocity expression under the conditions (*ε*, *χ*, *ε*) → (0,0,0) where 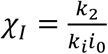.

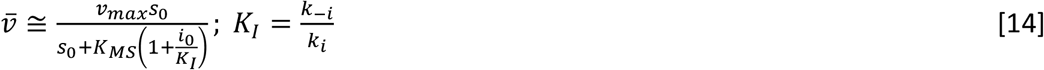

Similarly, for the pure uncompetitive inhibition given in Scheme C **of Fig. 1** one can derive the following average velocity expression.

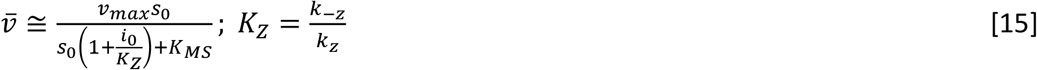

Finally, for the mixed competitive inhibition where both competitive and uncompetitive inhibition scenarios operate as given in Scheme C of **Fig. 1**, one can derive the following combined average velocity expression.

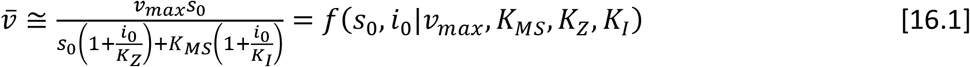

IC_50_ is the inhibitor concentration which is required to decrease 50% of the inhibitor free enzyme activity which can be straightforwardly calculated [23] as follows for the generalized mixed inhibition scenario by solving the following equation for i_0_.

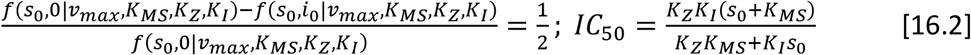

When the inhibition is non-competitive, then the inhibitor binds at the allosteric site of enzyme that is different from the substrate binding active-site. In such scenario, the average steady state velocity equation becomes as follows.

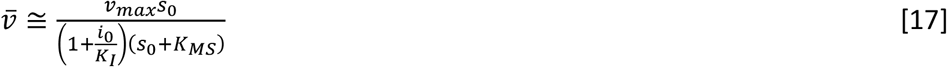

**Eq. 17** is similar to **Eq. 16.1** with *K*_*Z*_ = *K*_*I*_ and for non-competitive inhibition one finds that

*IC*_50_ = *K*_*I*_. For pure competitive inhibition one finds that 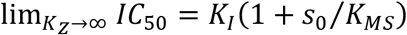 and for pure uncompetitive inhibition one finds that 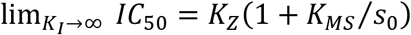 .

### Numerical Simulation

The stochastic version of nonlinear ODEs given in **Eqs. 1.1-1.4** can be numerically integrated using the following Euler type scheme on chemical Langevin equations [41, 42].

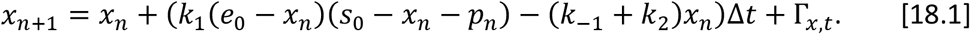

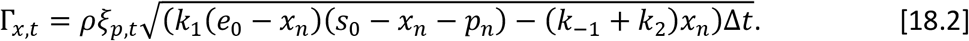

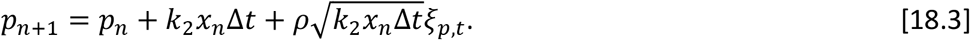

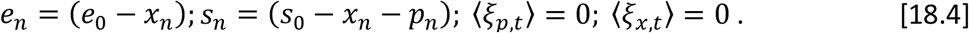

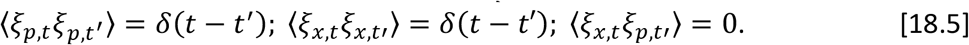

Here the initial conditions are *x* = 0, *s* = *s*_0_, *e* = *e*_0_, and *p* = 0 at *t* = 0, *ξ*_*q*,*d*_ where q = (x, p) are delta correlated gaussian white noise terms and *ρ* is the additional noise control parameter. The time increment parameter Δ*t* will be adjusted to capture both pre-as well as post-steady state dynamics. Setting *ρ* = 0, results in the deterministic trajectory of the MMS scheme for the given set of parameters and initial conditions. To check the consistency among the data fitting methods 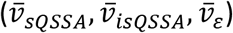, several stochastic trajectories of **Eqs. 1** were generated with fixed t_r_ and e_0_ and different s_0_ values. Stochastic simulated datasets are posted at (<u>DOI:</u> <u>10.5281/zenodo.15094623</u>) and the sample code implementations of ML algorithm are given in the **Supporting Materials**. To obtain the kinetic parameters k_1_, k_-2_ and k_2_ from the progress curve data with initial s_0_ and e_0_, we used nonlinear least square fitting [43] using Marquardt-Levenberg (ML) algorithm [44-47]. The computational workflow of ML scheme starts with minimizing the function 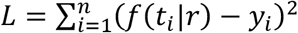.where *f*(*t*|*r*) is the numerical solution to **Eqs. 1** with the parameter set *r* = [*k*_1_, *k*_−1_, *k*_*2*_], and *y* is the observed dataset where (*f, y*) = (*p, s*). Upon expanding *L* in Taylor series around 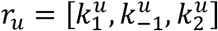 and then truncating with the first order terms, one finds the following approximation.

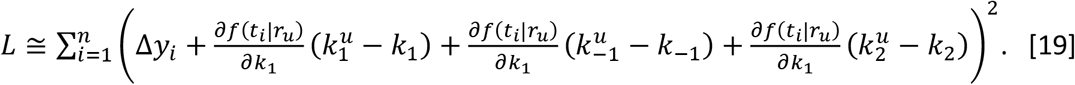

In this equation Δ*y*_*i*_ = *f*(*t*_*i*_ |*r*_*u*_) − *y*_*i*_ . The conditions for the minimum with respect to *r* will 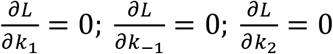. which can be explicitly written as follows.

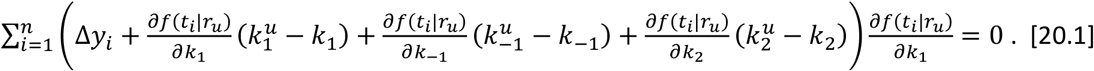

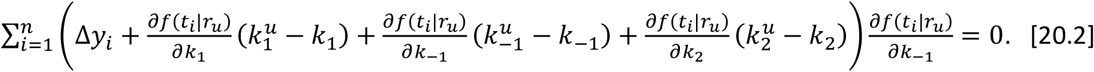

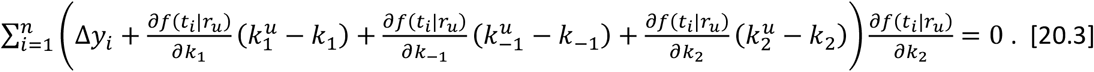

**Eqs. 15.1-3** can be written in a matrix form [43] as *J*^*T*^Δ*y* + (*J*^*T*^*J*)Δ*r* = 0. Definition of various terms in this matrix equation are given below.

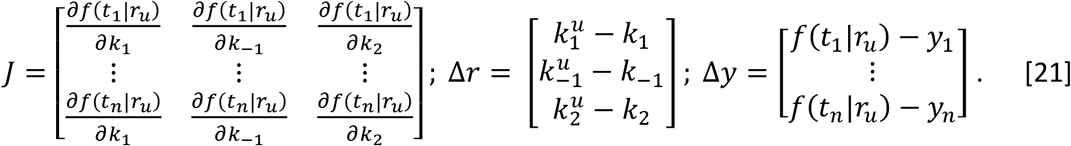

We compute the partial derivatives with respect to parameters [*k*_1_, *k*_−1_, *k*_*2*_] at time point t_i_ as follows where *f*(*t*|*r*) is the numerical integrated solution to **Eqs. 1** with the parameter set *r* = [*k*_1_, *k*_−1_, *k*_*2*_].

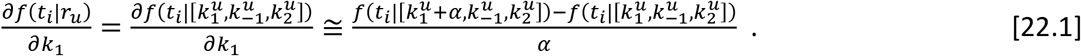

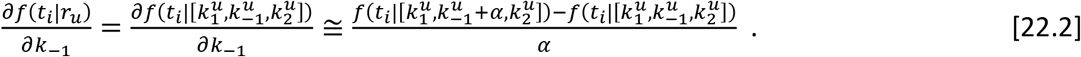

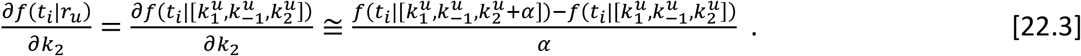

We set *α* to the lowest possible value depending on the computational accuracy requirement. In **Eqs. 16**, *J* is the Jacobian matrix and explicitly one finds the following.

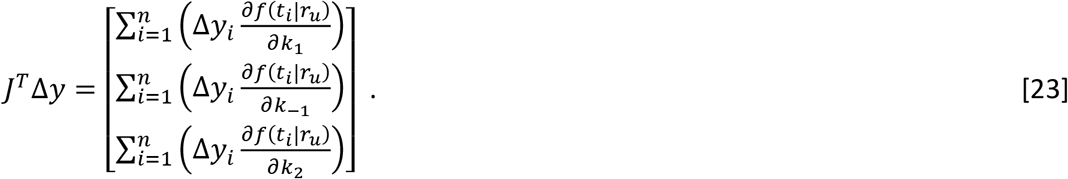

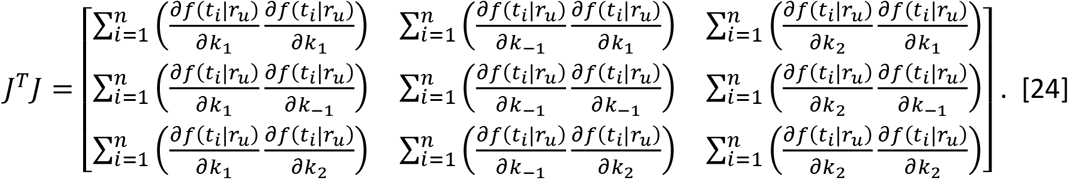

From **Eqs. 15-19**, one obtains the following Marquardt-Levenberg (ML) iterative scheme where λ is the ML parameter and λ = 0 represents the Newton’s method [43].

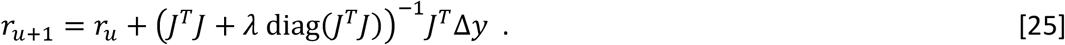

The reduced sum of squares of deviation will be computed at the end of u^th^ iteration as 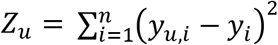. where *y*_*u*,*I*_ = *f*(*t*_*i*_ |*r*_*u*_) is the predicted data point values for *r*_*u*_ which will be obtained by numerical integration of **Eqs. 1** using Euler type scheme given in **Eqs. 13** with ρ = 0. Here λ should be adjusted depending on changing pattern of *Z*_*u*_ over iterations. When *Z*_*u*_ *< Z*_*u*+1_, then we set λ → λδ. When *Z*_*u*_ > *Z*_*u*+1_ then we set λ → λ⁄δ. Here the δ is the ML tuning parameter [44]. When |*Z*_*u*_ − *Z*_*u*+1_| *<* θ where θ is the required tolerance limit, then we exit from iteration and the final parameters 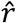 will be obtained. For NLS fit of various datasets we used λ = 10^−*2*^, δ = *2* and θ = 10^-6^. the final sum of squares error (SSE) and the covariance matrix will be calculated as follows.

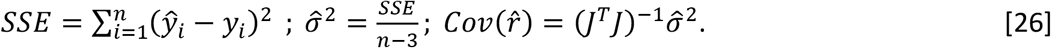

Upon computing the total sum of squares (SST), one can compute the final nonlinear regression coefficient R^2^ as follows.

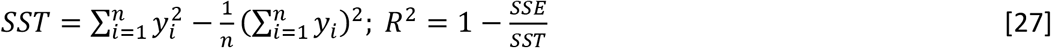

Here 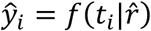 where 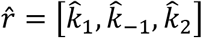 is the final fit parameter values. We use the final SSE and R^2^ to compare the goodness of fit across trajectories with different *s*_0_ and replications. Similar workflow was used to obtain K_M_ and v_max_ using 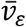 as given in **Eq. 10** with the median of k_1_ obtained from the progress curve analysis across different s_0_ over several replications outlined in **Eqs. 14-21**. Linear least square fitting with **Eq. 11** [43] for 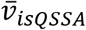 yields the required parameter estimates along with the errors.

### Multiple nonlinear regression for competitive inhibition

In this section, we will outline the multiple nonlinear regression fitting procedure for the Dixon type steady state average velocity inhibition dataset on 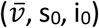, over **Eqs. 13-17**. For example, let us consider the case of pure competitive inhibition i.e., 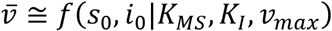as given in **Eq. 14**. For simplicity we use the notation (*a, b, c*) = (*K*_*MS*_, *K*_*I*_, *v*_*max*_) for the parameters and (*s*_0,*i*_, *i*_0,*k*_) = (*x*_*i*_, *y*_*k*_) for the concentrations of substrate and inhibitor. We denote the substrate levels *e*_0,*i*_ where i = 1 to n and inhibitor levels *i*_0,*k*_ where k = 1 to m and the corresponding average velocities as 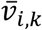 = product formed in the reaction time t_r_ / total reaction time t_r_. With these settings one can define the chi-square function to be minimized as follows.

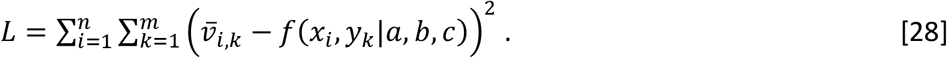

Here L needs to be minimized with respect to the parameters (a, b, c) along with two different independent random variables. In this equation, 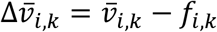 where *f*_*i*,*k*_ = *f*(*x*_*i*_, *y*_*k*_|*a*_0_, *b*_0_, *c*_0_) is the fit function. As in case of single variable nonlinear fitting procedure, we expand L in terms of Taylor’s series around (a_0_, b_0_, c_0_) and truncate the series with the first order term as follows.

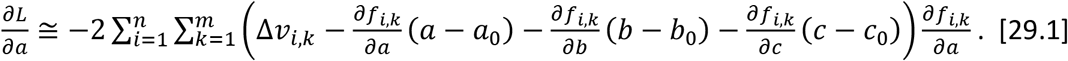

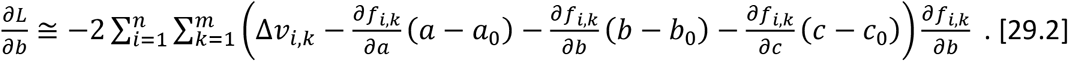

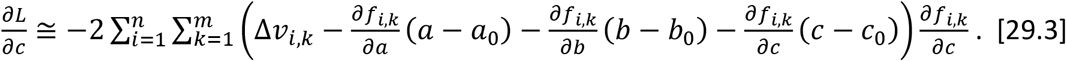

**Eqs. 24** can be su*m*arized in the following matrix form.

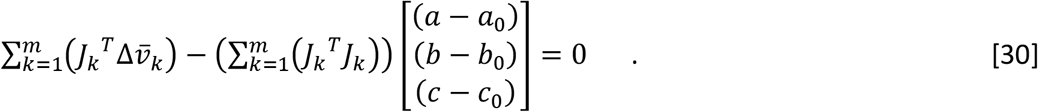

Various terms in **Eq. 25** are defined as follows.

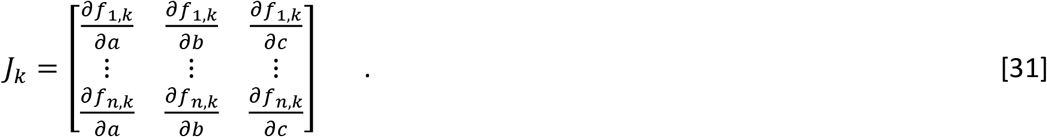

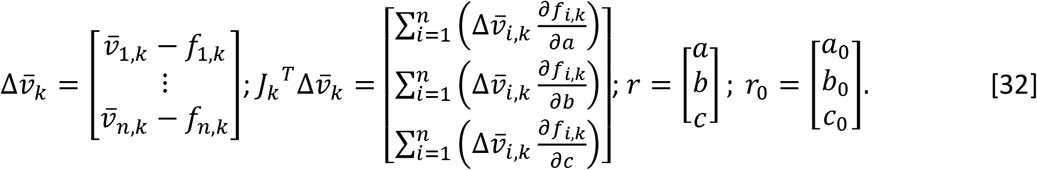

From **Eqs. 25**, one finds the Newton’s iterative scheme for the multiple nonlinear fitting procedure to minimize L as follows.

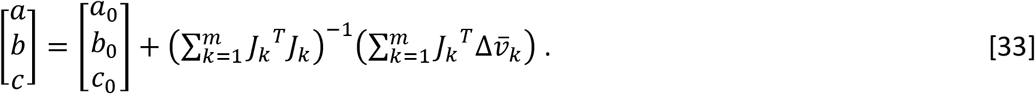

Clearly, before the inversion of J^T^J matrix, one needs to sum over the index k. Various partial derivative terms can be numerically computed as follows. This idea can be extended to any number of independent random variables apart from (s_0_ and i_0_).

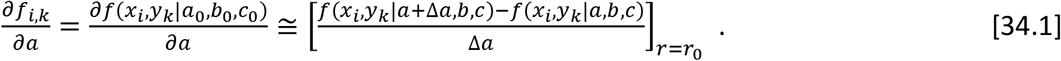

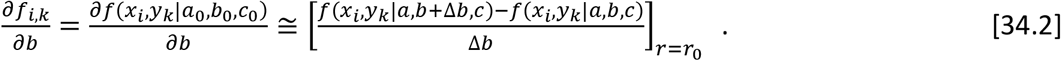

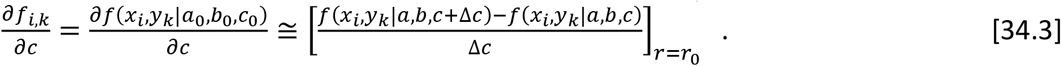

As in case of single variable NLS fitting described in **Eqs. 20**, Marquardt-Levenberg algorithm can be embedded in the **Eq. 26** to speed up the convergence. Sample datasets for competitive inhibitions schemes described by **Eqs. 13-17** were generated by adding gaussian white noise terms to the Dixon type steady state velocity datasets with appropriate noise control parameter *ρ* as 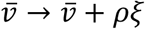 where *ξ* is the random number drawn from the standard normal population N (0, 1).

## Results and Discussion

There are several issues and limitations associated with the currently available steady state as well as progress curve methods to estimate the K_M_ and v_max_ of single substrate MM enzymes. First let us consider sQSSA with stationary reactant assumption 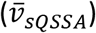 which is valid only when *t*_*c*_ ≪ *t*_*r*_ *< t*_*m*_ and (*η, ε*) → (0,0). Since K_M_ values are not generally available for the unknown enzyme systems, to achieve these criteria, one needs to set higher values for *s*_0_ than *e*_0_ which will lead to high background fluctuations in the measured average reaction velocities. For example, let 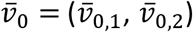 correspond to respectively *s*_0_ = (*s*_0,1_, *s*_0,*2*_). When *s*_0_ → ∞, then one finds the following.

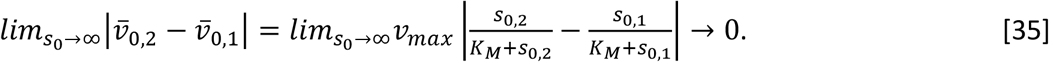

**Eq. 23** leads to high noise level in the observed average reaction velocities that in turn results in high level of uncertainty in the estimation of K_M_. Similar effects are prevalent in the progress curve analysis using especially the trajectories on (*p, t*) space since *pp* ≅ 0 in the pre-steady state regime that leads to high level of background fluctuations in the measured values of *p*. This in turn results in high uncertainty in the estimation of K_M_ using the data on (*p, t*) which significantly depends on the pre-steady state dynamics. Clearly, the data on (*p, t*) can be used only in the steady state methods which mainly work well in the post-steady state regime. Detailed NLS fitting analysis of the data on (*p, t*) and (*s, t*) spaces in the presence as well as absence of noise suggests the following.

a. Progress curve methods using Marquardt-Levenberg algorithm outlined in the methods section works very well over data on (s, t) as well as (p, t) in the absence of noise i.e. *ρ* = 0 in **Eqs. 13** as demonstrated in **Figs. 2A** and **2B**. NLS fitting over the progress curve data on (p, t) as well as (s, t) can recover the original K_M_ and v_max_ with minimal error.
b. When *ρ* > 0, then the NLS fit procedure work better with the data on (s, t) than the data on (p, t) as demonstrated in **Figs. 2C-F**. Particularly, fluctuations in the trajectories of (p, t) in the pre-steady state regime is demonstrated in **Fig. 2C**. NLS fitting methods can capture k_1_ and k_2_ very well and the estimate on k_-1_ is prone to severe uncertainties especially when k_-1_ ≪ (k_1_, k_2_) as demonstrated with sample trajectories from **Figs. 2C** and **2D** in **Figs. 2E** and **2F**. This in turn leads to significant amount error in the estimation of K_M_ although v_max_ is not much affected by this issue.
c. The distribution of NLS fit parameters obtained at different noise levels are shown in **Figs. 3** for both (s, t) and (p, t) time course datasets. These results clearly suggest that K_M_ values obtained from the NLS fit of time course dataset on (p, t) space are unreliable and prone to significant fluctuations in the presence of high noise levels. When the noise level increases, then the distribution of fit parameters corresponding to (p, t) space dataset exhibits bimodality as shown in (**Figs. 3D, 3F**). Whereas, NLS fit over time course dataset on (s, t) space can yield reliable median estimates of K_M_ even at high noise levels. Particularly, monomodal type distribution of fit values of K_M_ and v_max_ are observed for the dataset on (s, t) space as demonstrated in **Figs. 3A, 3C, 3E**.
d. Remarkably both time course datasets on (s, t) and (p, t) spaces can yield reliable NLS fit median estimates of v_max_ across various noise levels as demonstrated in **Fig. 3**. This observation is reasonable since v_max_ mostly relies on the post-steady state dynamics and therefore the fluctuations in the pre steady state regime of (p, t) space will not influence the error in the estimation of v_max_.

**FIGURE 2.**
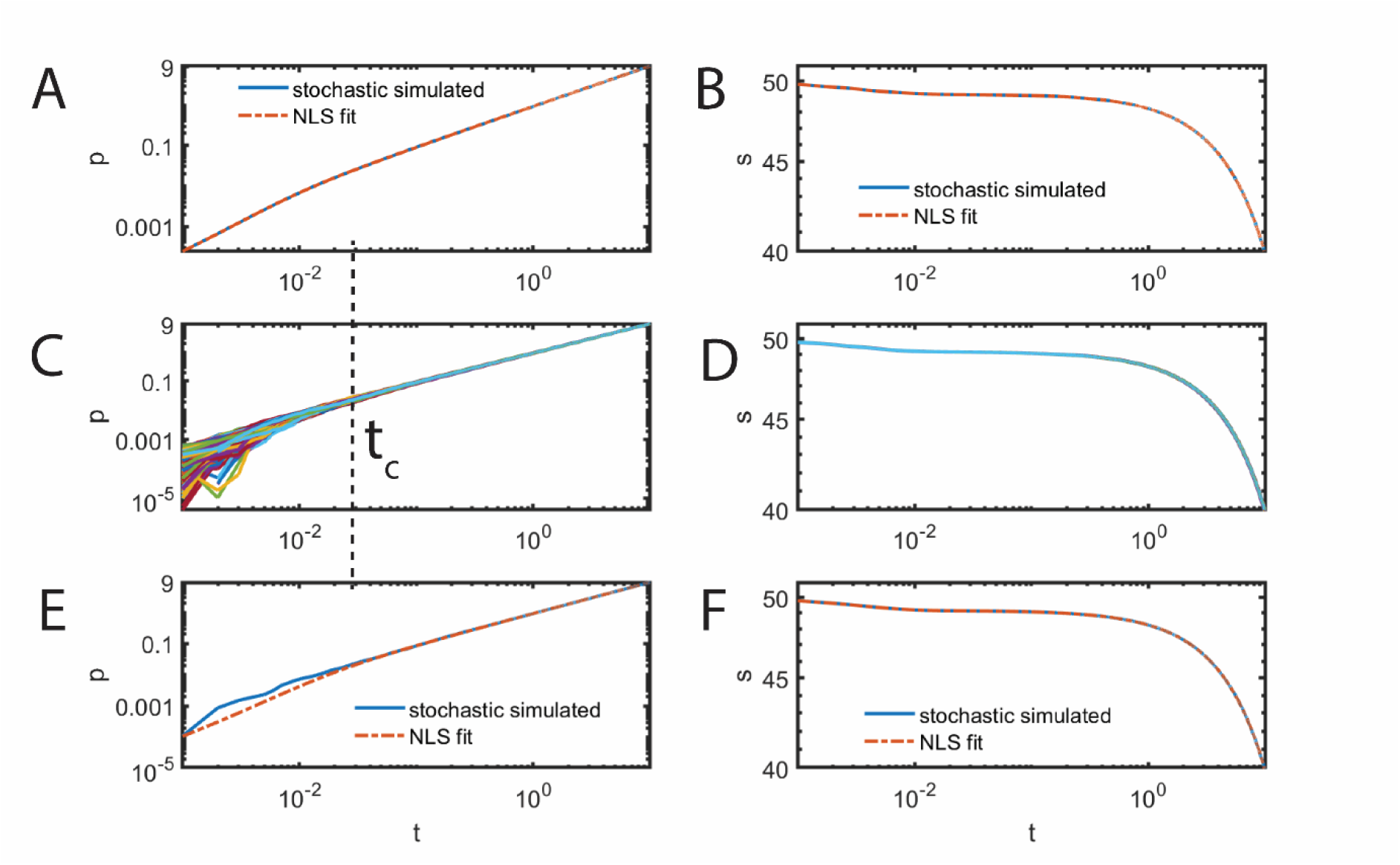
Progress curve analysis of data on (p, t) and (s, t). Co*m*on simulation settings for the numerical integration scheme in **Eqs. 13** are k_1_ = 5.1 µM^-1^s^-1^, k_-1_ = 0.5 s^-1^, k_2_ = 1.1 s^-1^, Δ*t* = 10^-3^ s, e_0_ = 0.85 µM, s_0_ = 50 µM so that K_M_ = 0.313725 µM, t_c_ ≈ 0.0039 s, t_m_ ≈ 118.38 s, total simulation time = 10 s and v_max_ = 0.935 µM s^-1^. **A**. Simulation of **Eqs. 13** with *ρ* = 0 and NLS fit of the simulation trajectory with **Eqs. 1** using ML algorithm yielded the parameters k_1_ = 5.1 ± 1.25 x 10^-5^, k_-1_ = 0.499 ± 7 x 10^-6^, k_2_ = 1.1 ± 4.5 x 10^-8^ for the data on (p, t) with K_M_ = 0.314 at 95% confidence level. **B**. NLS fit using ML algorithm over **Eqs. 1** yielded k_1_ = 5.1 ± 6 x 10^-6^, k-1 = 0.499 ± 9 x 10^-6^, k_2_ = 1.1 ± 4.5 x 10^-8^ for data on (s, t) generated using ρ = 0 in **Eqs. 13** with K_M_ = 0.313639 at 95% confidence level. **C**. stochastic trajectories (1000 numbers) of **Eqs. 13** with ρ = 10^-2^ demonstrating the fluctuations in the pre-steady state regime of (p, t) space. **D**. stochastic trajectories of **Eqs. 13** with ρ = 10^-2^ demonstrating less fluctuations in the pre-steady state regime of (s, t) space. **E**. NLS fit of sample (p, t) trajectory with ρ = 10^-2^ yielded k_1_ = 2.29± 0.09, k_-1_ = 1.53 ± 0.1, k_2_ = 1.12 ± 2 x 10^-3^ with K_M_ = 1.18 and v_max_ = 0.95 at 95% confidence level. **F**. NLS fit of sample (s, t) trajectory with ρ = 10^-2^ over **Eqs. 1** yielded k_1_ = 5.1 ± 0.07, k_-1_ = ± 0.1, k_2_ = 1.1 ± 5 x 10^-4^ with K_M_ = 0.47 and v_max_ = 0.94 at 95% confidence level.

**FIGURE 3.**
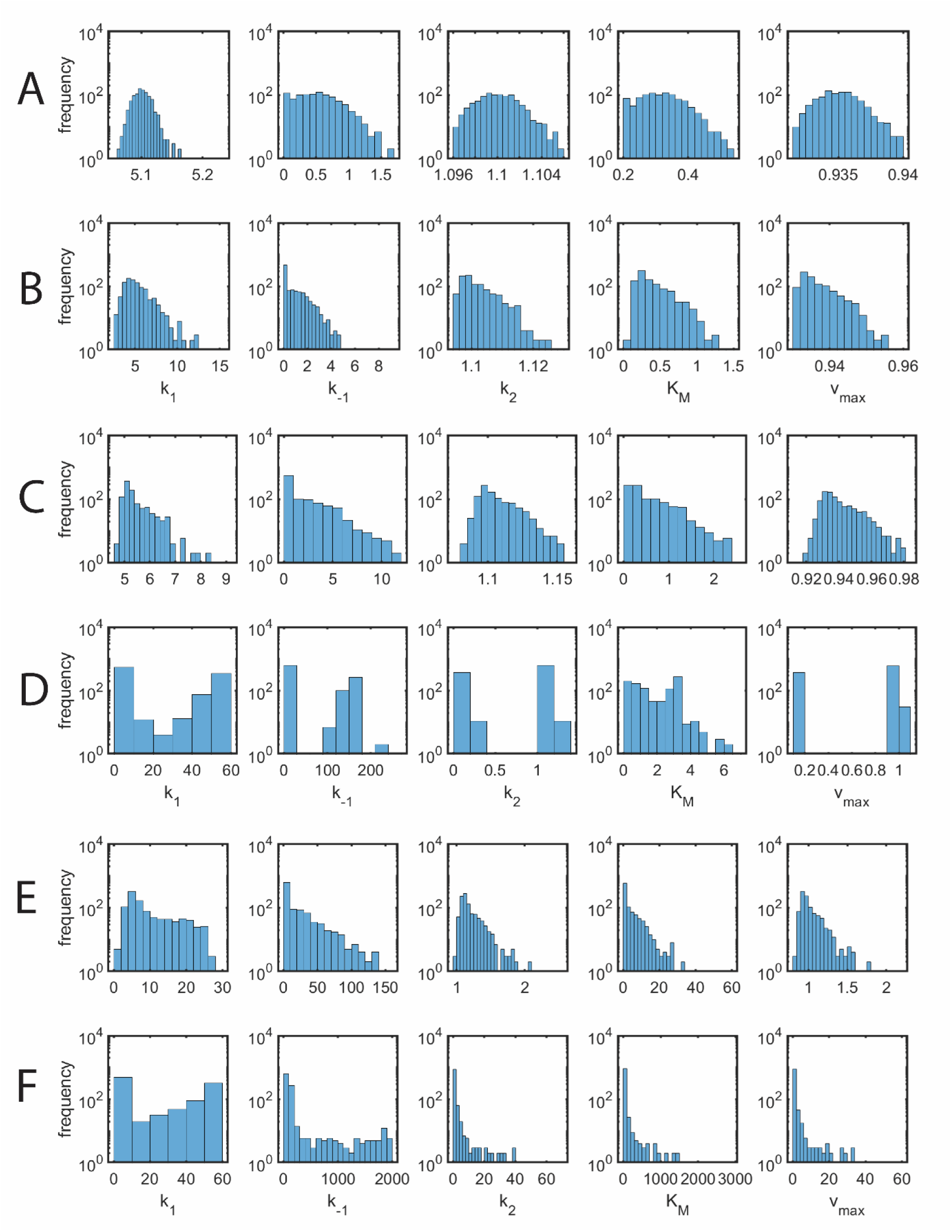
Distribution of nonlinear least squares (NLS) fit parameters across various noise levels. Co*m*on simulation settings for the numerical integration scheme in **Eqs. 13** are k_1_ = 5.1 µM^-1^s^-1^, k_-1_ = 0.5 s^-1^, k_2_ = 1.1 s^-1^, Δ*t* = 10^-3^ s, total simulation time = 10 s, *e*_0_ = 0.85 µM so that K_M_ = 0.313725 µM and v_max_ = 0.935 µM s^-1^, s_0_ = 50 µM for which one finds that t_c_ ≈ 0.0039 s and t_m_ ≈ 118.38 s. NLS fit was done using Marquardt-Levenberg algorithm as outlined in the methods section. In A, C and E, NLS fittings were done using data on (s, t). Whereas, in B, D and F the fittings were done using data on (p, t). Upon obtaining k_1_, k_-1_ and k_2_, K_M_ and v_max_ will be calculated. Noise levels in **Eqs. 13** were set as follows. **A, B**. ρ = 10^-3^. **C, D**. ρ = 10^-2^. **E, F**. ρ = 10^-1^. The median values of K_M_ and v_max_ were computed across 10^3^ replications. **A**. the median values of (k_1_, k_-1_, k_2_, K_M_, v_max_) = (5.1005, 0.5009, 1.100, 0.3140, 0.9350). **B**. the median values of (k_1_, k_-1_, k_2_, K_M_, v_max_) = (4.9188, 0.3849, 1.1001, 0.3163, 0.9351). **C**. the median values of (k_1_, k_-1_, k_2_, K_M_, v_max_) = (5.2031, 0.5499, 1.1021, 0.3197, 0.9368). **D**. the median values of (k_1_, k_-1_, k_2_, K_M_, v_max_) = (4.7469, 1.1193, 1.0981, 1.5888, 0.9334). **E**. the median values of (k_1_, k_-1_, k_2_, K_M_, v_max_) = (6.5060, 0.4776, 1.1348, 0.3727, 0.9646). **F**. the median values of (k_1_, k_-1_, k_2_, K_M_, v_max_) = (16.0345, 0.7332, 1.1380, 3.1860, 0.9673).

These observations suggest that neither progress curve analysis alone nor steady state methods alone can provide consistent estimates of K_M_. Multiple ways of analyzing the same dataset of (p, t) and (s, t) is clearly required to check for the consistency across various steady state and progress curve methods and obtain reliable estimates. In this context, one can use the median value of k_1_ obtained from progress curve methods into the NLS fit on 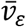 as given in **Eq. 10** to obtain consistent estimates of K_M_ and v_max_ across progress curve analysis and steady state methods. We consider the following methods I, II, III and IV using the same main time course datasets on (s, p, t). For Methods II, III and IV, sub-dataset on average reaction velocity versus *s*_0_ at different t_r_ will be generated from the main dataset. Here the average velocity is defined as (total product formed at the end of t_r_) / t_r_ where t_r_ will be iterated approximately from the steady state timescale t_c_ to t_m_. Various NLS fit methods I, II, III and IV are su*m*arized as follows.

a. **Method I** is the direct NLS fitting of the main progress curve dataset on (s, t) with fixed *e*_0_ and different *s*_0_ across several replications with the original rate equations **Eqs. 1.1-1.4** using Marquardt-Levenberg algorithm as outlined in the methods section. K_M_ values will be calculated from the fit values of (k_1_, k_-1_, k_2_) and the median of (k_1_, k_-1_, k_2_, K_M_) values will also be obtained.
b. In **Method II**, the parameter k_1_ in the definition of 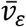 of **Eqs. 10** will be fixed with the median of k_1_ values obtained from Method I and the sub-dataset on 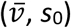, computed from the main time course dataset on (p, t) at different t_r_ will be used for NLS fitting with fixed k_1_ to obtain K_M_ and v_max_.
c. In **Method III**, 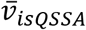 as defined in **Eq. 11** will be used to fit the sub-dataset on 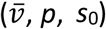 computed from the main dataset on (p, t) at different t_r_ and s_0_. One can also use 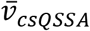 that is computationally costlier than 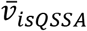
d. In **Method IV** that is widely used across various literature, 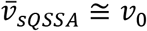 will be used to fit the sub-dataset on 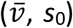 computed from the main dataset on (p, t) at different reaction time t_r_ and s_0_.

In Methods II, III and IV, the K_M_ value corresponding to t_r_ with minimum sum of square error (SSE) as given in **Eq. 21** will be computed as demonstrated in **Fig. 4**. There exists a critical reaction time t_r_ at which the sum of square error in the estimation of K_M_ attains the minimum which is clearly observed in the absence of noise in the dataset (ρ = 0, **Fig. 4A**). Remarkably, **Fig. 4B** suggest that the error in the estimation of K_M_ strongly depends on the reaction time t_r_ in steady state Methods II, III and IV. In the presence of noise in the dataset, the median of SSE corresponding to K_M_ decreases monotonically and the fit parameter value converges to the original K_M_ as t_r_ is increased towards t_m_. The speed of convergence of fit K_M_ towards the original K_M_ is much faster in Method II than in Methods III and IV as demonstrated in **Fig. 4B**. Sum of square errors of various approximations can be defined as 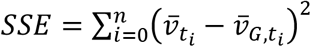 where *t*_*n*_ = *t*_*r*_ and 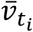 is the numerically simulated average velocity at time t = t_i_, and 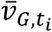 are the approximations where the subscripts G = sQSSA, csQSSA, ε, isQSSA (I, II) associated with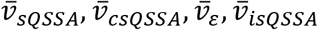. The computed *SSE* seems to be strongly dependent on t_r_ and range of iteration of s_0_ as demonstrated in **Fig. 4C** and **4D**. Clearly, SSE of is minimum near t_c_ and the SSEs corresponding to 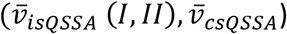 attain minimum approximately near t_m_. The timescale t_Q_ at which the SSE of 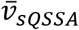 attains minimum can be computed by solving 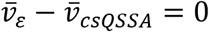 for t as shown in **Fig. 4E**. This is the critical time point at which both pre- and post-steady state approximations 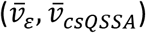 start to deviate from the original simulated average reaction velocity values.

**FIGURE 4.**
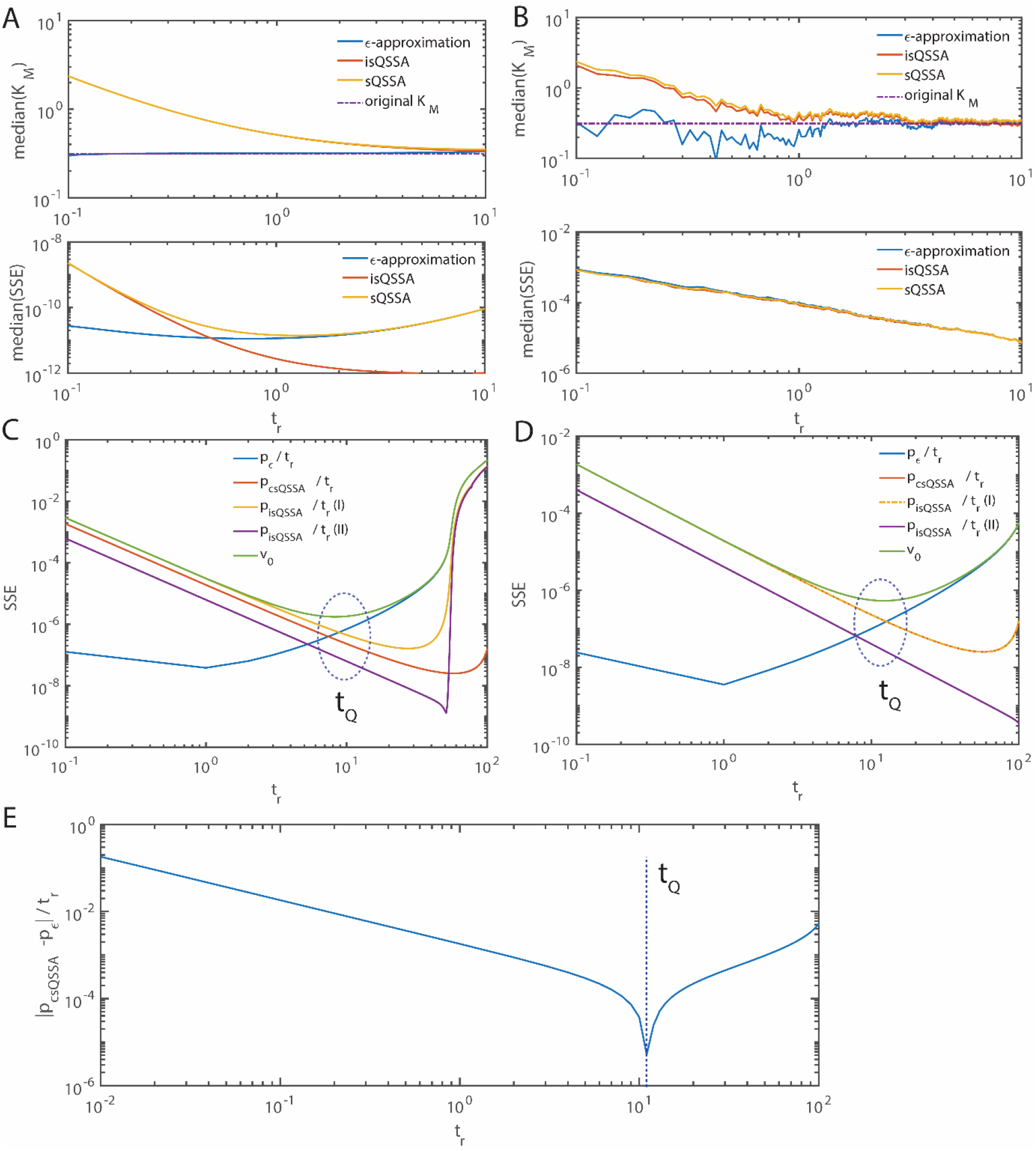
Variation of the fitted K_M_ values with respect to reaction time t_r_ across different fitting methods. Co*m*on simulation settings for the numerical integration scheme in **Eqs. 13** are k_1_ = 5.1 µM^-1^ s^-1^, k_-1_ = 0.5 s^-1^, k_2_ = 1.1 s^-1^, Δ*t* = 10^-3^ s, total simulation time = 10 s, *e*_0_ = 0.85 µM so that K_M_ = 0.313725 µM and *v*_*max*_ = 0.935 µM s^-1^, s_0_ was iterated from 50 to 250 µM with interval of 25 µM. For *s*_0_ = 50 µM one finds that *t*_*c*_ ≈ 0.0039 s and *t*_*m*_ ≈ 118.38 s. The reaction time t_r_ was iterated from 0.1 to 10 s. The sub datasets on 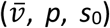, at different t_r_ were generated from the main progress curve dataset on (p, t) and used to fit **Eqs. 10** (ε-approximation), **Eq. 11** (isQSSA) and v_0_ (sQSSA) and the median values of K_M_ were obtained across 10^3^ replications. **A**. ρ = 0 in **Eqs. 13. B**. ρ = 10^-2^ in **Eqs. 13. C**. Δ*t* = 10^-5^ s, s_0_ was iterated from 50 to 250 µM with interval of 25 µM. For p_csQSSA_ / t_r_, s_0_ was iterated from 100 to 200 µM with interval of 10 µM. **D**. For all the steady state methods, Δ*t* = 10^-5^ s, s_0_ was iterated from 100 to 200 µM with interval of 10 µM. **E**. s_0_ = 100 µM and Δ*t* = 10^-5^ s for which t_Q_ ≈ 10.49 s.

When *t*_*r*_ = *t*_*Q*_, then the reaction velocity will be 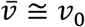 leading to minimum amount of error in the approximation 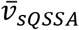 Remarkably, all the fitting methods 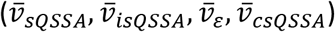 approximately perform well near t_Q_, at which the variation of the fit parameters across these methods will be at minimum. The entire procedure will be repeated with different *e*_0_ until obtaining consistent median estimates of K_M_ and v_max_ with minimal threshold variation across methods I to IV. We consider the coefficient of variation (CV = standard deviation / mean) of the median K_M_ values obtained from methods I-IV as the *reliability index*. **Figs. 4C, 4D** and **4E** suggest that the reliability index will be at minimum when the reaction time is set as t_r_ = t_Q_. In the absence of *a priori* knowledge on enzyme kinetic parameters, one can use the reliability index to obtain best fit values of K_M_ which occurs near the optimum reaction time t_Q_. Upon obtaining such consistent median estimates of K_M_ corresponding to a given threshold *reliability index*, all the K_M_ values obtained from the same dataset using different methods will be then pooled and the overall median of K_M_ and v_max_ will be computed as depicted in **Fig. 5**.

**FIGURE 5.**
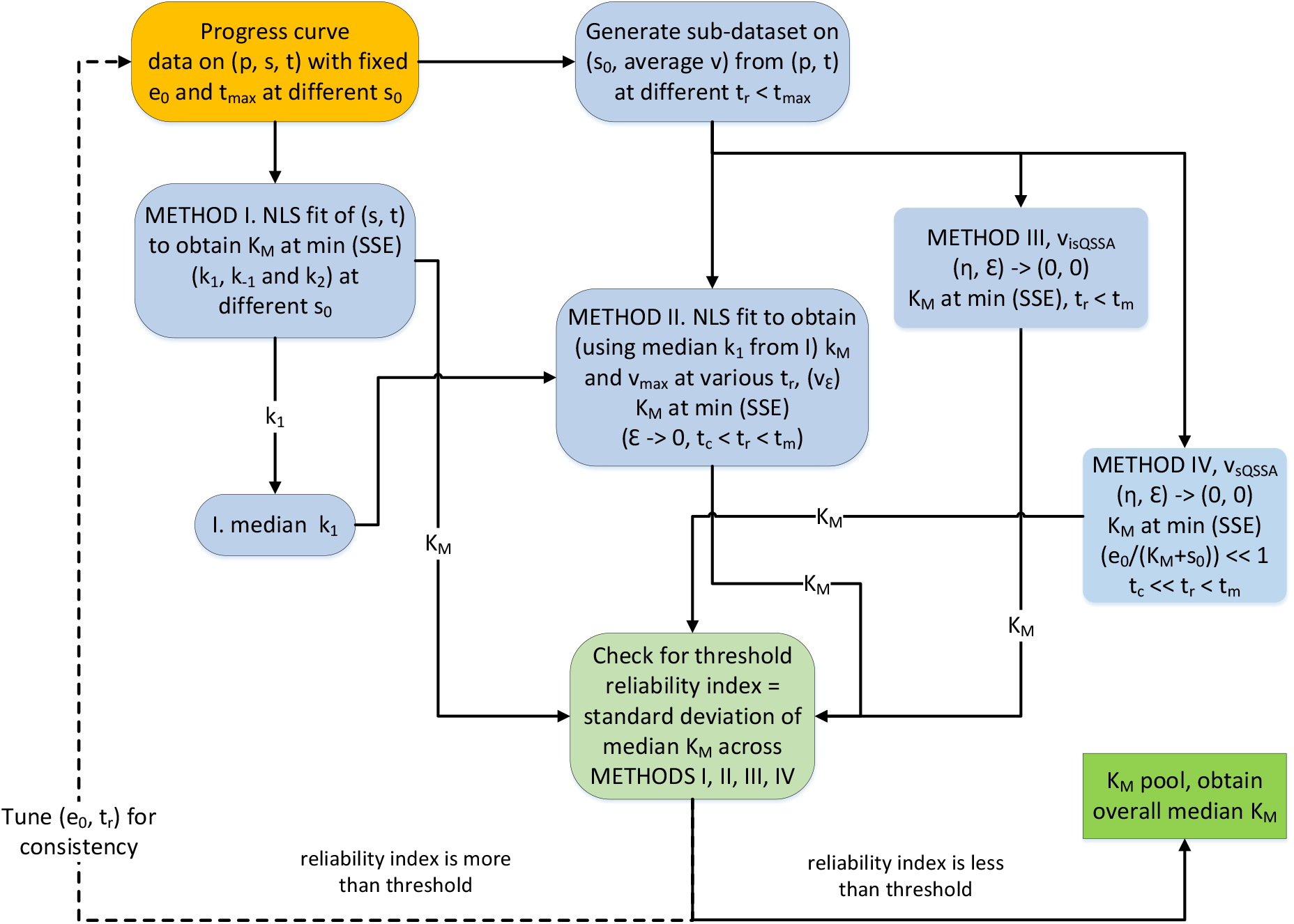
Computational algorithm to obtain consistent estimate of K_M_ from the progress curve dataset. Here, the progress curve data on (p, s, t) will be the main input for Methods I, II, III and IV. Here t_max_ is the maximum data collection time. Method I is the direct nonlinear least square fitting of the dataset on (s, t) with fixed *e*_0_ and different *s*_0_ across replications with the original **Eqs. 1.1-1.4** using Marquardt-Levenberg algorithm as outlined in the methods section. The parameter k_1_ in 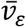 of **Eqs. 10** of Method II will be fixed with the median of k_1_ values obtained from Method I. For Methods II-IV, sub-dataset on average velocity ((totalproduct formed at the end of t_r_) / t_r_, where t_r_ will be iterated from t_c_ towards t_m_) versus *s*_0_ at different t_r_ will be generated from the main dataset. NLS fittings over 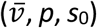, were done using models corresponding to Methods II-IV at different t_r_. The K_M_ value corresponding to t_r_ with minimum sum of square error (SSE) will be considered. Upon obtaining consistency on the estimates of K_M_ across Methods I-IV corresponding to the given threshold *reliability index*, all the K_M_ values obtained from the same dataset using different methods will be then pooled and the overall median K_M_ will be computed. Here *reliability index* is the coefficient of variation of median K_M_ values obtained from Methods I-IV.

The distributions of various fit parameters obtained from Methods I to IV over several replications are shown in **Fig. 6** along with the overall pooled distribution of K_M_ and v_max_. The distribution of fit parameters (k_1_, k_-1_, k_2_) show skewness towards lower values (**Figs. 6A1-3**). Particularly, the distribution of k_-1_ exhibits a bimodal type with zero spike. As result, the distribution of K_M_ also exhibits spike near zero and v_max_ exhibits skewed distribution (**Figs. 6B1-2**). Since the median of k_1_ obtained from Method I is used for the NLS fitting calculations in Method II, the distribution of K_M_ values obtained from Method II also exhibits a spike near zero as shown in **Fig. 6B-3**. Interestingly, both K_M_ and v_max_ obtained from Method III exhibit sy*m*etric type distribution (**Figs. 6C-1, 6D-1**). However, the distribution of K_M_ values obtained using Method III span towards negative values (**Fig. 6C-1**). The distribution of K_M_ values obtained from Method IV also exhibit a spike near zero (**Fig. 6C-2**). Clearly, the inconsistency among the K_M_ values obtained from different methods originates mainly from the type of distribution of fit parameter in the presence of noise.

**FIGURE 6.**
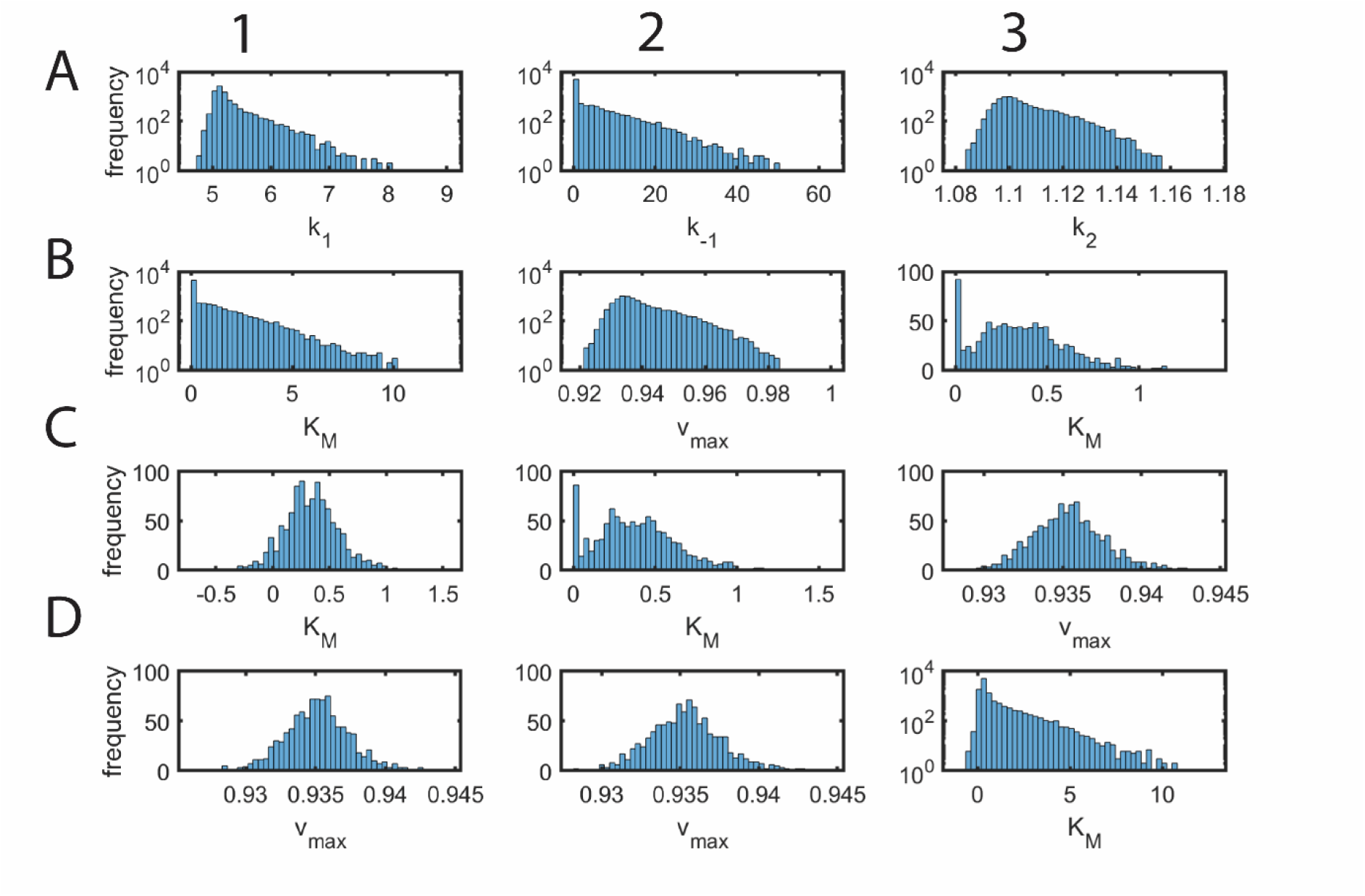
Distribution of fit parameters across various methods I-IV. Co*m*on simulation settings for the numerical integration scheme in **Eqs. 13** are k_1_ = 5.1 µM^-1^s^-1^, k_-1_ = 0.5 s^-1^, k_2_ = 1.1 s^-1^, Δ*t* = 10^-3^ s, total simulation time = 10 s, *e*_0_ = 0.85 µM so that K_M_ = 0.313725 µM and v_max_ = 0.935 µM s^-1^, s_0_ was iterated from 50 to 250 µM with interval of 25 µM. For *s*_0_ = 50 µM one finds that t_c_ ≈ 0.0039 s and t_m_ ≈ 118.38 s. The median values corresponding to progress curve fit over 10^3^ trajectories on (s, t) dataset were (k_1_, k_-1_, k_2_, K_M_, v_max_) = (5.164, 0.587, 1.102, 0.331, 0.937). **A1-3, B1-2**. NLS fit method I using Marquardt-Levenberg algorithm. Upon obtaining k_1_, k_-1_ and k_2_, K_M_ and v_max_ will be calculated as in **B1, B2. B3, C3**. Method II. Here the obtained median values from datasets on (p, t) were (K_M_, v_max_) = (0.338, 0.935). **C1, D1**. Method III. Here the obtained median values from datasets on (p, t) were (K_M_, v_max_) = (0.332, 0.935). **C2, D2**. Method IV. Here the obtained median values from datasets on (p, t) were (K_M_, v_max_) = (0.366, 0.935). **D3**. Overall distribution of pooled K_M_ values across all the methods I-IV with overall median of K_M_ = 0.3418 µM which correspond to the coefficient of variation of median K_M_ values obtained from Methods I, II, III and IV as 0.048.

Efficient progress curve fitting also seems to be dependent on the detailed data collection around the transition regions in (s, t) and (p, t) spaces with maximum change in curvature [37]. Clearly, there exists at least three different timescale regimes in the trajectory on (p, t) space with significant change in curvature viz. exponential increasing phase in the pre-steady state regime, linear increasing phase in the steady state region and reaction-ending phase. Similar timescale regimes also exist in (s, t) space. Significant change in the curvature occurs at the interface of these timescale regimes approximately near t_c_ and t_m_. Accurate values of these interface timescales can be calculated from *e*_*ε*_ and *e*_*csQSSA*_ for the pre- and post-steady state regimes respectively. The approximate curvature *ζ*_*d*_ associated with *e*_*ε*_ of **Eq. 8** can be defined as follows [48].

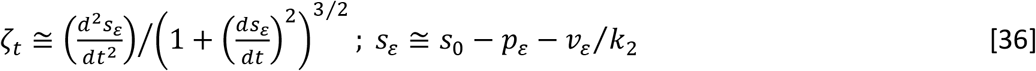

Here *p*_*ε*_ and *v*_*ε*_ are defined as in **Eqs. 8**. Upon solving 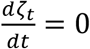 for t, one finds the time scale associated with the maximum change in curvature at the interface of pre-steady to steady

steady-state transition *t*_*R*_ as follows.

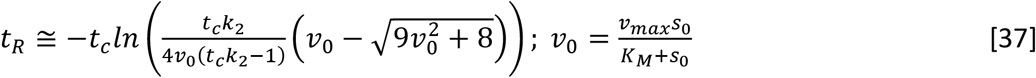

Similarly noting the definition of *e*_*csQSSA*_ as given in **Eqs. 12**, the approximate curvature *ψ*_*t*_ in the steady state to post-steady state transition regime of (s, t) space can be defined as follows.

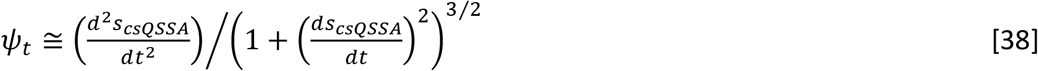

Upon solving 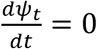 for t, one finds the timescale associated with the maximum change in curvature at the interface of steady state to reaction ending regime *t*_*U*_ as follows.

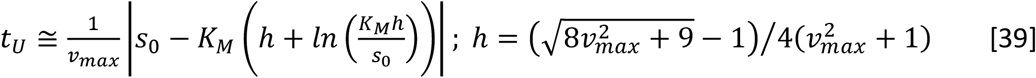

Since the enzyme system evolves with constant velocity *v*_0_ in the timescale region *t*_*R*_ *< t < t*_*U*_, progress curve fitting methods can yield reliable estimate of K_M_ only when the experimental data collection is done around the timescales (t_R_, t_U_) with maximum change in the curvature of (s, t) trajectory. Whereas, steady state methods work very well in the constant velocity regime with minimal change in the curvature of the trajectory of (s, t). Change in curvature of the trajectory in (s, t) space with respect to time is shown in **Fig. 7A** along with the approximations given in **Eqs. 25-27** which are in good agreement with the numerical simulation results. Further, **Eqs. 25** and **27** suggest that (t_R_, t_U_) will be close to (t_c_, t_m_) and for practical purposes one can use (t_c_, t_m_) for checking the validity conditions of various model fitting methods. The simulated average reaction velocity (p / t) along with the approximations 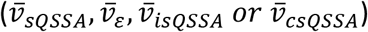 are shown in **Fig. 7B**. These suggest that the approximation 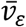 works well in the pre-steady to steady state regime and 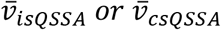 work well in the post steady state regime and 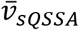 work only in narrow range of timescales with constant reaction velocity. Clearly, the reaction time t_r_ plays important role in deciding the reliability index among Methods I to IV. The error in the estimation of K_M_ using 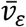 proportionally increases when t_r_ increases well beyond the steady state. Conversely, inclusion of data points from the pre-steady state regime increases the error in 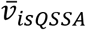 or 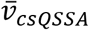 These means that the error in the estimation of KM using the widely used 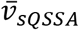 will be always higher than the error associated with 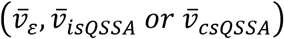. Reliability index can be used as a check point to optimize the appropriate t_r_ to obtain accurate estimate of K_M_ values from progress curve and various steady state methods.

**FIGURE 7.**
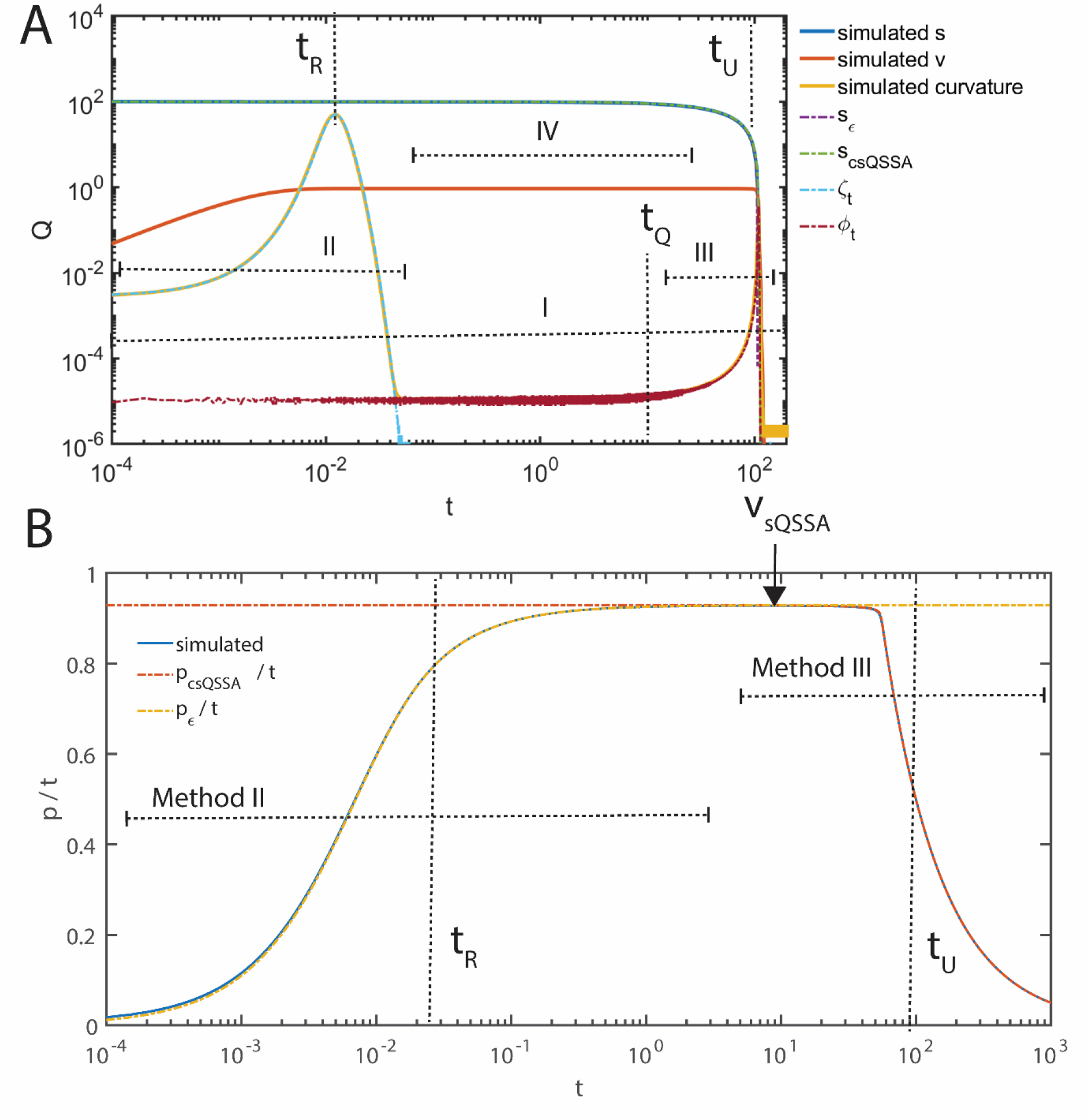
Change in the curvature of trajectory on (s, t) space and average reaction velocity. Co*m*on simulation settings for the numerical integration scheme in **Eqs. 13** are k_1_ = 5.1 µM^-1^ s^-1^, k_-1_ = 0.5 s^-1^, k_2_ = 1.1 s^-1^, Δ*t* = 0.5 x 10^-4^ s, total simulation time = 1000 s, noise level set as ρ = 0, *e*_0_ = 0.85 µM so that K_M_ = 0.313725 µM and *v*_*max*_ = 0.935 µM s^-1^, and s_0_ = 100 µM. For these settings, one finds that *t*_*c*_ ≈ 0.002 s, *t*_*m*_ ≈ 91.2 s and *t*_*Q*_ ≈ 10.49 s. **A**. The transition timescales (t_R_, t_U_) at which maximum change in the curvature occurs can be calculated using **Eqs. 25-27** as t_R_ = 0.0126 s and t_U_ = 92 s. The time dependent curvatures associated with the trajectory in (s, t) space *ζ*_*d*_ and *ψψ*_*d*_ corresponding to the pre- and post-steady state regimes respectively are defined as in **Eqs. 24-27**. Progress curve Method I works very well in the regions with maximum change in the curvature. Whereas, steady state methods II, III and IV work very well in the regions with minimal change in the curvature. **B**. Change in the average velocity (p / t) with time. Method II works well from the pre-steady to steady state regime. Whereas, Method III works well in the steady to post-steady state regime. Method IV works well only in a narrow timescale regime with constant average velocity.

Software packages such as global kinetic explorer and dynafit [49-51] were developed to fit the data obtained from progress curve and steady state experiments. Several comparative studies were done earlier over steady state and progress curve methods to optimize the analysis workflow towards minimizing the estimation error [52, 53]. Selective removal of data points [54] over certain progress curve regime were also proposed. However, all these methods are solely either progress curve or steady state type and there is no way to check the consistency of fit parameters across various methods since each method requires different type of input dataset. Accurate estimation of kinetic parameters depends on strict impose of validity conditions of each method as given in **Table 1**. Unfortunately, these validity conditions require *a priori* knowledge on various enzyme kinetic parameters which are not generally available for unknown systems. Since the conditions of validity for each method is different, a global analysis workflow with a comon input dataset is required to obtain consistent estimates. In this context, our proposed computational workflow depicted in **Fig. 5** can yield reliable estimates of K_M_ from the comon time course dataset at different initial substrate concentrations s_0_ and replications. The reliability index which is the coefficient of variation of median K_M_ values obtained from Methods I-IV can act as a check point against unreliable estimates of K_M_. When the computed reliability index is more than the given threshold value, then one needs to iterate the reaction time until obtaining the desired threshold level. Clearly, the proposed methods do not require *a priori* knowledge on K_M_ values unlike the currently available sQSSA methods.

**TABLE 1.** Conditions of validity of various fitting methods to obtain K_M_ and v_max_.

| Description | Definition | Validity conditions |
| --- | --- | --- |
| Progress curve analysis | Time course dataset on (s, p, t). | timescale interval should capture both pre- and post-steady state regimes. (Method I) |
| $\bar{v}_{sQSSA}$ | $\bar{v}_{sQSSA} \cong v_{max}s_0/(K_M + s_0)$ | $t_c \ll t_r < t_m, (\eta, \varepsilon) \rightarrow (0,0), (e_0/(K_M + s_0)) \ll 1$, Method IV |
| $\bar{v}_{isQSSA}$ | $\bar{v}_{isQSSA} \cong v_{max} + \frac{K_M}{t_r} \ln\left(\frac{s_0 - p}{s_0}\right)$ (I)<br>$\bar{v}_{isQSSA} \cong v_{max} + \frac{K_M}{t_r} \ln\left(\frac{s}{s_0}\right)$ (II) | $t_c < t_r < t_m, (\eta, \varepsilon) \rightarrow (0,0)$<br>Method III. |
| $\bar{v}_{csQSSA}$ | $\bar{v}_{csQSSA} \cong \frac{1}{t_r}(s_0 - s_{csQSSA})$<br>$s_{csQSSA} \cong K_M W\left(\frac{s_0}{K_M} \exp\left(\frac{s_0 - v_{max}t_r}{K_M}\right)\right)$ | $t_c < t_r < t_m, (\eta, \varepsilon) \rightarrow (0,0)$<br>Method III. Alternate form of $\bar{v}_{isQSSA}$. |
| $\bar{v}_\varepsilon$ | $\bar{v}_\varepsilon \cong v_0(1 - t_c h(t_r)/t_r)$<br>$v_0 = v_{max}s_0/(K_M + s_0)$<br>$t_c \cong 1/k_1(K_M + s_0)$<br>$h(t_r) = (1 - \exp(-t_r/t_c))$ | $t_r < t_m, \varepsilon \rightarrow 0$<br>Method II. median of $k_1$ obtained from Method I will be substituted in $t_c$. |

Identification of the correct inhibition mode using the experimental dataset on 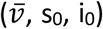 viz. competitive, uncompetitive, noncompetitive and mixed is essential to design drug molecules for the target enzyme and obtain their IC_50_ values for the comparison purposes. Dixon type graphical methods [26] used to identify the type of inhibition are not accurate in the presence of noise in the dataset on 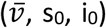. It will be difficult to visually identify and measure the intersection points in the Dixon plot. Sequential and simultaneous nonlinear regression methods [27, 28] were proposed to overcome such issues. In this context, we have developed the multiple nonlinear regression (MNLS) method to evaluate various inhibition parameters directly by fitting dataset on 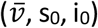 over the respective inhibition model equations as shown in **Fig. 8**. Using MNLS method, one can directly obtain the parameters fit values (K_M_, K_I_, K_Z_, v_max_) along with their standard errors from the Dixon type datasets via minimizing the overall sum of square deviations as defined in **Eq. 28**. Fit parameter values for sample datasets in the presence and absence of noise are given in **Tables 2** and **3**. MNLS method can recover the exact parameter values in the absence of noise in the datasets as demonstrated in **Figs. 8A, 8C, 8E, 8F** and **Table 2**. Further, an algorithm based on the sum of square error (SSE) values of various multiple nonlinear fit functions and standard errors of fit parameters is also developed as shown in **Fig. 9**. In this algorithm, the given enzyme inhibition dataset on 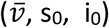 will be simultaneously fitted with competitive, uncompetitive, noncompetitive and mixed inhibition model equations as given in **Eqs. 14-17**. The overall sum of square errors computed with respect to each of these model functions were compared and the model function showing the lowest overall SSE will be chosen as the best fit model. When the chosen model is a mixed-competitive inhibition model, then further investigations will be carried out as follows to avoid ambiguities arising due to the generalized nature of mixed inhibition model equation.

**TABLE 2.** Multiple nonlinear regression of enzyme inhibition data in the absence of noise.

| Parameter fit | Competitive | Uncompetitive | Non competitive | Mixed |
| --- | --- | --- | --- | --- |
| Competitive inhibition dataset |  |  |  |  |
| $K_M$ | $2.81 \pm 1.4 \times 10^{-10}$ | $709019 \pm 0.3$ | $141.9 \pm 7.2$ | $2.81 \pm 5.4 \times 10^{-9}$ |
| $K_I$ | $0.355 \pm 1.7 \times 10^{-11}$ | - | $9.48 \pm 0.4$ | $0.355 \pm 7 \times 10^{-10}$ |
| $K_Z$ | - | $0.00124 \pm 3.6 \times 10^{-10}$ | - | $278930 \pm 15$ |
| $v_{max}$ | $4.62 \pm 8.3 \times 10^{-12}$ | $35824.9 \pm 1.4 \times 10^{-10}$ | $13.21 \pm 0.47$ | $4.62 \pm 6.1 \times 10^{-9}$ |
| SSE | 9.64496e-25 | 0.0220512 | 0.00144751 | 1.08116e-11 |
| Uncompetitive inhibition dataset |  |  |  |  |
| $K_M$ | $0.003 \pm 1.6 \times 10^{-9}$ | $2.80952 \pm 8.8 \times 10^{-10}$ | $0.36 \pm 0.03$ | $2.81 \pm 7.5 \times 10^{-7}$ |
| $K_I$ | $0.0005 \pm 2.6 \times 10^{-10}$ | - | $2.3 \pm 0.029$ | $118254 \pm 103$ |
| $K_Z$ | - | $2.12903 \pm 1.2 \times 10^{-9}$ | - | $2.13 \pm 8.8 \times 10^{-9}$ |
| $v_{max}$ | $1.4 \pm 5 \times 10^{-7}$ | $4.62 \pm 2.3 \times 10^{-10}$ | $4.32072 \pm 0.047$ | $4.62 \pm 1.8 \times 10^{-8}$ |
| SSE | 0.0159053 | 2.29059e-23 | 1.59847e-06 | 2.80365e-14 |
| Noncompetitive inhibition dataset |  |  |  |  |
| $K_M$ | $0.00184 \pm 6 \times 10^{-5}$ | $135276 \pm 382$ | $2.81 \pm 4.1 \times 10^{-10}$ | $2.81 \pm 8.8 \times 10^{-9}$ |
| $K_I$ | $0.000189 \pm 6 \times 10^{-6}$ | - | $0.35 \pm 2.4 \times 10^{-10}$ | $0.35 \pm 1 \times 10^{-9}$ |
| $K_Z$ | - | $0.00021 \pm 4.3 \times 10^{-8}$ | - | $0.35 \pm 9 \times 10^{-11}$ |
| $v_{max}$ | $0.33 \pm 0.007$ | $7454.34 \pm 1.4$ | $4.62 \pm 3 \times 10^{-9}$ | $4.62 \pm 1.2 \times 10^{-9}$ |
| SSE | 0.000445374 | 2.17735e-06 | 4.79886e-24 | 2.81224e-25 |
| Mixed inhibition dataset |  |  |  |  |
| $K_M$ | $8.4 \times 10^{-5} \pm 2 \times 10^{-3}$ | $383529 \pm 287$ | $14.985 \pm 0.14$ | $2.81 \pm 2 \times 10^{-9}$ |
| $K_I$ | $7.8 \times 10^{-6} \pm 10^{-4}$ | - | $1.7 \pm 0.04$ | $0.35 \pm 3 \times 10^{-10}$ |
| $K_Z$ | - | $0.0004 \pm 7 \times 10^{-8}$ | - | $2.13 \pm 1 \times 10^{-10}$ |
| $v_{max}$ | $1.58 \pm 2.39$ | $20448 \pm 3$ | $5.625 \pm 0.1$ | $4.62 \pm 2 \times 10^{-9}$ |
| SSE | 0.0038 | 0.000377 | $1.63 \times 10^{-6}$ | $2 \times 10^{-24}$ |

**TABLE 3.** Multiple nonlinear regression of enzyme inhibition data in the presence of noise.

| Parameter fit | Competitive | Uncompetitive | Non competitive | Mixed |
| --- | --- | --- | --- | --- |
| Competitive inhibition dataset |  |  |  |  |
| $K_M$ | $2.87 \pm 0.14$ | $642519 \pm 0.06$ | $142.1 \pm 16.7$ | $2.73 \pm 3.9$ |
| $K_I$ | $0.3 \pm 0.02$ | - | $9.5 \pm 0.6$ | $1.17 \pm 31$ |
| $K_Z$ | - | $0.0014 \pm 8.6 \times 10^{-11}$ | - | $5.6e+06 \pm 0$ |
| $V_{max}$ | $4.62 \pm 0.009$ | $32449 \pm 2 \times 10^{-3}$ | $13.3 \pm 12.2$ | $4.70554 \pm 3.9$ |
| SSE | $1.063e-06$ | 0.0220808 | 0.00145037 | 0.830483 |
| Uncompetitive inhibition dataset |  |  |  |  |
| $K_M$ | $0.0031 \pm 3 \times 10^{-7}$ | $2.81 \pm 0.19$ | $0.37 \pm 0.04$ | $2.67 \pm 0.2$ |
| $K_I$ | $0.0005 \pm 5 \times 10^{-8}$ | - | $2.28 \pm 0.04$ | $243.82 \pm 388$ |
| $K_Z$ | - | $2.12 \pm 0.03$ | - | $2.12532 \pm 0.004$ |
| $V_{max}$ | $1.4 \pm 9 \times 10^{-5}$ | $4.66 \pm 0.05$ | $4.34 \pm 0.06$ | $4.62 \pm 0.008$ |
| SSE | 0.0159049 | $1.06653e-06$ | $2.477e-06$ | $1.07078e-06$ |
| Noncompetitive inhibition dataset |  |  |  |  |
| $K_M$ | $0.002 \pm 8.3 \times 10^{-6}$ | $104307 \pm 409$ | $2.78 \pm 0.19$ | $0.0023 \pm 0.001$ |
| $K_I$ | $0.0002 \pm 9 \times 10^{-7}$ | - | $0.35 \pm 0.11$ | $0.0003 \pm 0.0001$ |
| $K_Z$ | - | $0.00026 \pm 7 \times 10^{-8}$ | - | $0.38 \pm 0.02$ |
| $V_{max}$ | $0.33 \pm 9 \times 10^{-4}$ | $5928.66 \pm 1.58$ | $4.67 \pm 1.43$ | $4.314 \pm 0.17$ |
| SSE | 0.00044 | $3.51923e-06$ | $1.01839e-06$ | $1.01705e-06$ |
| Mixed inhibition dataset |  |  |  |  |
| $K_M$ | $6.2e-05 \pm 1 \times 10^{-9}$ | $329020 \pm 42$ | $14.95 \pm 0.19$ | $1.89 \pm 1.3$ |
| $K_I$ | $5.7e-06 \pm 1 \times 10^{-10}$ | - | $1.7 \pm 0.05$ | $0.24 \pm 0.16$ |
| $K_Z$ | - | $0.00045 \pm 1 \times 10^{-8}$ | - | $2.16 \pm 0.06$ |
| $V_{max}$ | $1.58 \pm 2 \times 10^{-5}$ | $17591 \pm 0.45$ | $5.63 \pm 0.14$ | $4.55 \pm 0.12$ |
| SSE | 0.00381337 | 0.000379777 | $2.76054e-06$ | $8.3677e-07$ |

**FIGURE 8.**
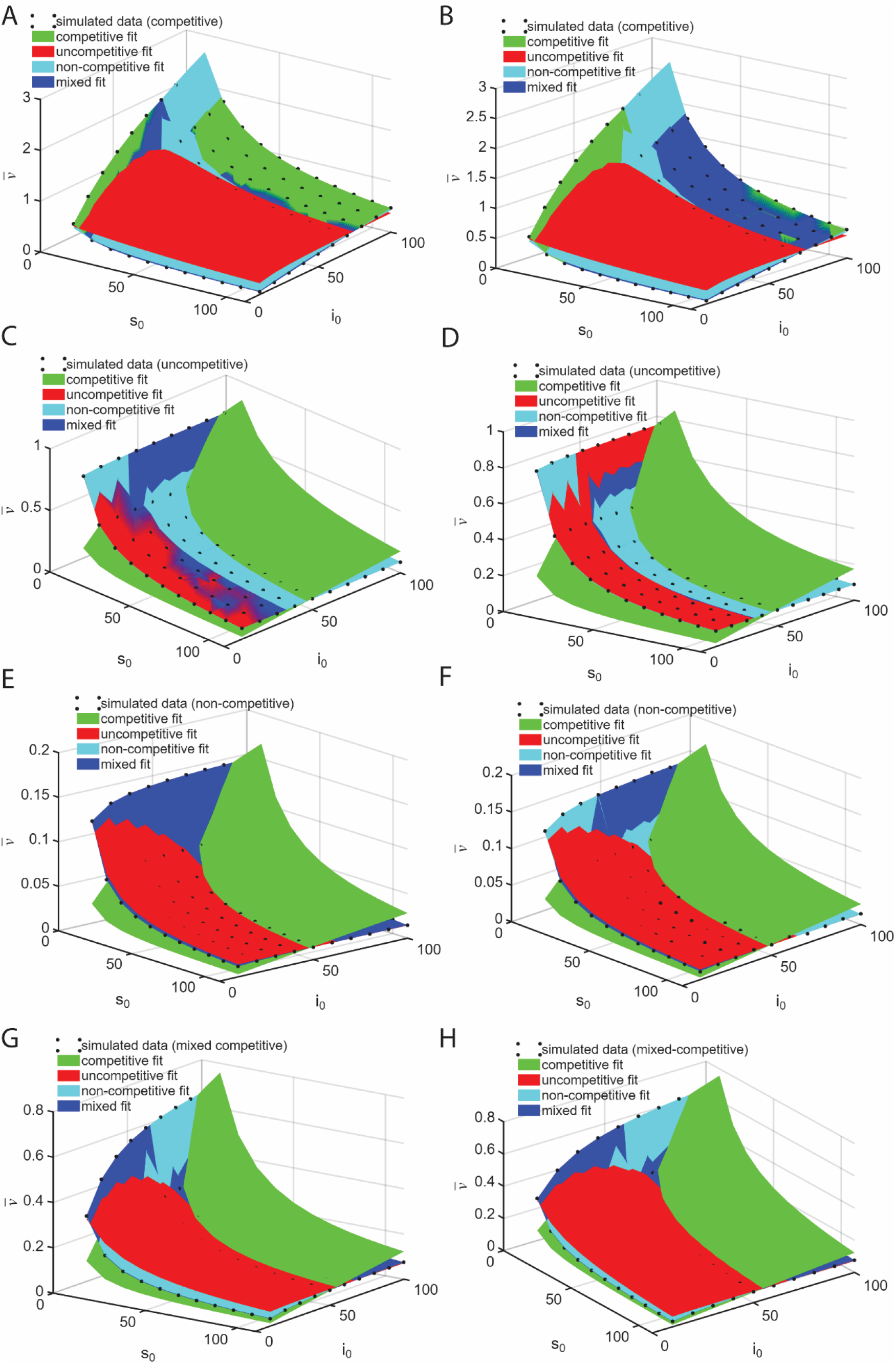
Multiple nonlinear regression fitting of Dixon type enzyme inhibition dataset on 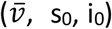, in the presence and absence of noise. A, C, E and G are in the absence of noise. B, D, F and H are in the presence of noise with strength ρ = 10^-3^. The preset values of parameters are K_M_ = 2.80952 µM, K_I_ = 0.354839 µM, K_Z_ = 2.12903 µM and v_max_ = 4.62 µM/s. s_0_ was iterated from 10 to 100 µM with an increment of 10 µM and i_0_ was iterated in 10 to 110 µM with an increment of 10 µM. **Eqs. 14-17** were used to generate datasets for (1) competitive, (2) uncompetitive, (3) noncompetitive, (4) mixed-competitive modes of enzyme inhibition and the multiple nonlinear fitting were done as outline in the methods section. The fit results are su*m*arized **Tables 2** and **3. A, B**. Competitive inhibition data fitted with (1)-(4) modes of inhibition. **C, D**. Uncompetitive inhibition data fitted with (1)-(4) modes of inhibition. **E, F**. Noncompetitive inhibition data fitted with (1)-(4) modes of inhibition. **G, H**. Mixed-competitive inhibition data fitted with (1)-(4) modes of inhibition.

**FIGURE 9.**
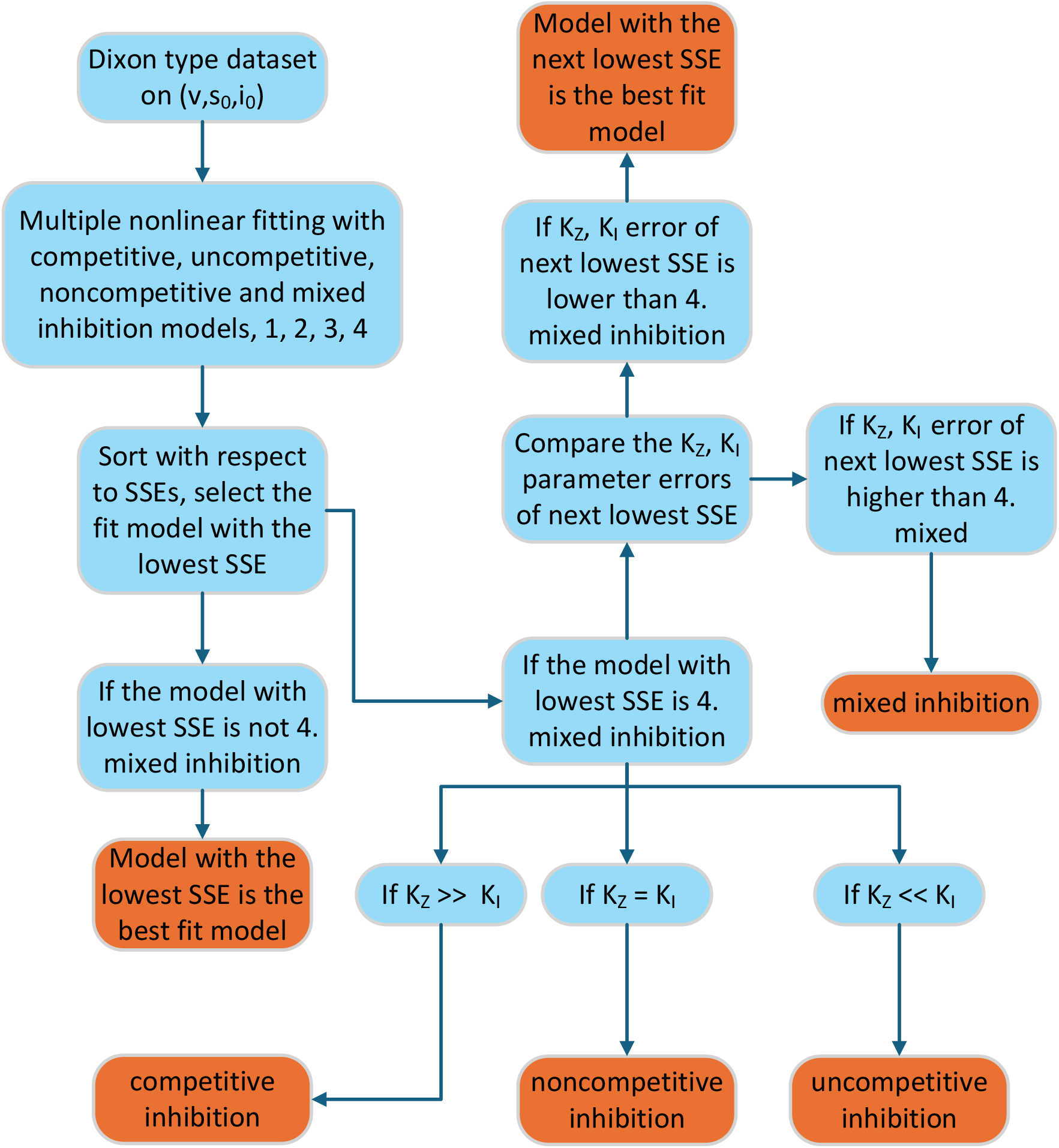
Classification algorithm for the type of inhibition from Dixon type dataset. Here the given dataset on 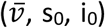 was fitted with competitive, uncompetitive, noncompetitive and mixed inhibition models and the sum of square errors were compared and model showing lowest overall SSE will be chosen. If the model is a mixed-competitive inhibition model, then the following investigations should be carried out. (1) when K_I_ = K_Z_, then it will the noncompetitive inhibition model. (2) when the parameter error corresponding to K_I_ or K_Z_ is higher than the next lowest SSE, then the next lowest SSE model will be the best fit model. (3) when K_I_ >> K_Z_, then it will be the uncompetitive inhibition model. (4) when K_I_ << K_Z_, then it will be the competitive inhibition model.

a. When K_I_ = K_Z_, then it will be interpreted as the noncompetitive inhibition model. This scenario occurs when one tries to fit noninhibition dataset over mixed inhibition model function.
b. When the parameter error corresponding to K_I_ or K_Z_ is higher than the next lowest SSE, then the next lowest SSE model will be the best fit model.
c. When K_I_ >> K_Z_, then it will be interpreted as the uncompetitive inhibition model. When the dataset on uncompetitive inhibition is fitted with mixed inhibition model equation, then one observes K_I_ >> K_Z_ in the fit results as shown in **Tables 2** and **3**.
d. When K_I_ << K_Z_, then it will be interpreted as the competitive inhibition model. When the dataset on competitive inhibition is fitted with mixed inhibition model equation, then one observes K_I_ << K_Z_ in the fit results as shown in **Table 2** and **3**.

Performance of the inhibition type classification algorithm in the presence of noise is shown in **Fig. 10**. Clearly, our inhibition type classification algorithm works very well especially at low noise levels. Further, fitting of competitive inhibition dataset may be wrongly shown as mixed inhibition type at moderate noise levels as demonstrated in **Fig. 10A**. At high noise levels, uncompetitive inhibition dataset may be wrongly shown as (**Fig. 10B**) noncompetitive inhibition and noncompetitive inhibition dataset may be wrongly detected as mixed, competitive and uncompetitive inhibition (**Fig. 10C**). These results clearly suggests that repeated data collection and MNLS analysis is required to obtain consistent classification of inhibition schemes and the respective parameters and IC_50_.

**FIGURE 10.**
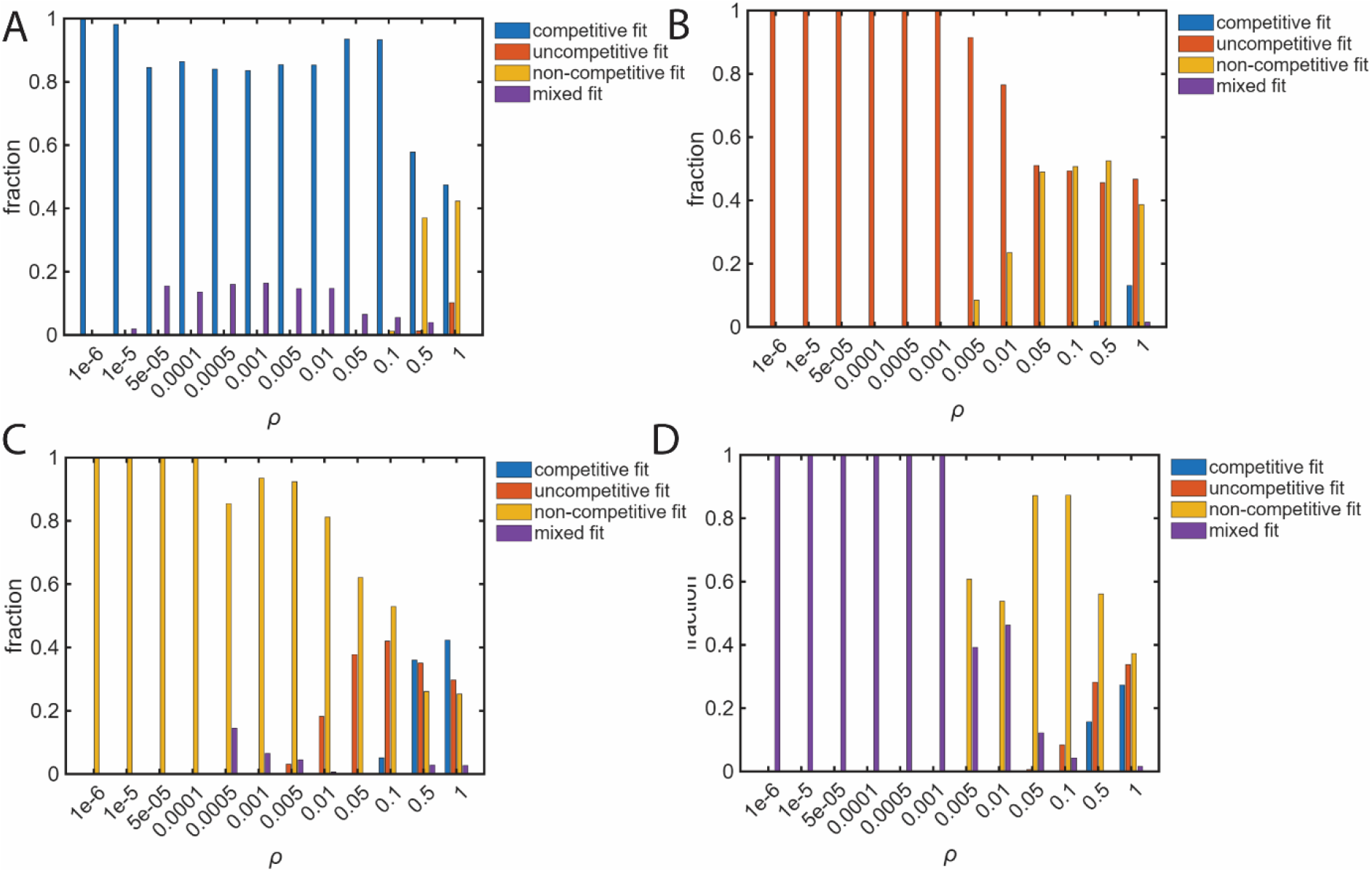
Performance of the inhibition type classification algorithm. The preset values of parameters are K_M_ = 2.80952 µM, K_I_ = 0.354839 µM, K_Z_ = 2.12903 µM and v_max_ = 4.62 µM/s. s_0_ was iterated from 10 to 100 µM with an increment of 10 µM and i_0_ was iterated in 10 to 110 µM with an increment of 10 µM. **Eqs. 14-17** were used to generate datasets for (1) competitive, (2) uncompetitive, (3) noncompetitive, (4) mixed-competitive modes of enzyme inhibition and the multiple nonlinear fitting were done as outlined in the methods section. Gaussian white noise with zero mean and unit variance was added through the control parameter ρ that was iterated from 10^-6^ to 1 as 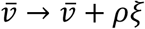 where *ξ* is the random number drawn from the standard normal population N (0, 1). Fraction fit results were computed over 1000 randomly generated datasets. **A**. Competitive inhibition datasets were fitted with model equations corresponding to (1)-(4) modes of inhibition. **B**. Uncompetitive inhibition datasets were fitted with model equations corresponding to (1)-(4) modes of inhibition. **C**. Noncompetitive inhibition datasets were fitted with model equations corresponding to (1)-(4) modes of inhibition. **D**. Mixed-competitive inhibition datasets were fitted with the model equations corresponding to (1)-(4) modes of inhibition.

## Conclusion

Accurate estimation of enzyme kinetic parameters such as K_M_ and v_max_ from the experimental datasets is critical to characterize any enzyme. The validity of the widely used standard quasi steady state method over decades relies on the *a priori* information of K_M_ and other kinetic parameters which are generally not available for newly identified enzyme systems. Progress curve methods were proposed to alleviate such issues. However, nonlinear least square fitting of the time course dataset with the original differential rate equations works well only in the timescale regimes with maximum change in the curvature of the trajectory of substrate evolution. Identifying these timescales requires the knowledge on enzyme kinetic parameters. Though experimental trial and error methods can be used to identify such timescales, they are generally cost ineffective.

Further, progress curve methods work better with the time course data on substrate than the time course data on product since the product concentration is close to zero in the pre-steady state regime. Since steady state and progress curve analysis use different types of datasets in the current analysis set up, the estimated K_M_ values will be inconsistent among these methods. This leads to the dilema of identifying correct K_M_ values among various steady state and progress curve analyses. Here we have shown that the error in the estimation of K_M_ using steady state methods is strongly dependent on the total reaction time. We have further shown that the steady state methods work well over the reaction timescales with minimum change in the curvature of the trajectory of product evolution with constant reaction velocity. Considering these theoretical factors, we have developed an integrated method of analysis which comprises of both progress curve and steady state analysis using the same time course datasets that also consider the total reaction time. We considered three different methods of steady state analyses that involve fitting of the average reaction velocity with the total substrate concertation. We have defined the reliability index which is the coefficient of variation of the median K_M_ values obtained from progress curve and other steady state fitting methods using the same time course dataset at different substrate concentrations and replications. We further demonstrate that reliable estimate of K_M_ can be obtained by iterating the total reaction time to minimize the reliability index across time course and steady state curve fitting methods.

We have developed a multiple nonlinear fitting procedure-based algorithm to classify the average velocity, total substrate and inhibitor concentrations dataset over competitive, uncompetitive, noncompetitive and mixed types. This method works very well at low to medium noise levels. However, at high noise levels the competitive inhibition dataset may be wrongly shown as mixed inhibition type, uncompetitive inhibition dataset may be wrongly shown as noncompetitive inhibition and noncompetitive inhibition dataset may be wrongly detected as mixed, competitive and uncompetitive inhibition types. Repeated data collection and multiple nonlinear fitting analysis is required to obtain consistent estimates of inhibition parameters and IC_50_.

## Data availability statement

The datasets generated during and/or analysed during the current study are available in the Zenodo. DOI: 10.5281/zenodo.15094623

## SUPPORTING MATERIALS

### Datasets for the global analysis

DOI: 10.5281/zenodo.15094623

#### FIGURES 2C, 2D

sampleptr.txt and samplestr.txt

Simulation settings **(Eqs. 13**) are k_1_ = 5.1 µM^-1^s^-1^, k_-1_ = 0.5 s^-1^, k_2_ = 1.1 s^-1^, Δ*t*= 10^-3^ s, e_0_ = 0.85 µM, s_0_ = 50 µM and ρ = 10^-2^. Totally there are 1000 replications. First column is the time in seconds, and subsequent columns are concentrations of product (sampleptr.txt) and substrate (samplestr.txt) respectively.

#### FIGURE 3

Simulation settings (Eqs. 13) are k_1_ = 5.1 µM^-1^s^-1^, k_-1_ = 0.5 s^-1^, k_2_ = 1.1 s^-1^, Δ*t* = 10^-3^ s, e_0_ = 0.85 µM and s_0_ = 50 µM. Totally there are 1000 replications. First column is the time in seconds, and subsequent columns are replications of concentrations of product (sampleptr.txt) and substrate (samplestr.txt) respectively.

sampledat_ptpm1.txt, sampledat_ptpm2.txt, sampledat_ptpm3.txt respectively with ρ = 10^-1^, 10^-2^ and 10^-3^ for product evolution.

sampledat_stpm1.txt, sampledat_stpm2.txt, sampledat_stpm3.txt respectively with ρ = 10^-1^, 10^-2^ and 10^-3^ for substrate evolution.

#### FIGURES 2E, 2F, 4 and 6

Datasets were generated by numerical integration of **Eqs. 13** as given in main text with the following settings k_1_ = 5.1 µM^-1^s^-1^, k_-1_ = 0.5 s^-1^, k_2_ = 1.1 s^-1^, ρ = 10^-2^, Δ*t* = 10^-3^ s, e_0_ = 0.85 µM, s_0_ was iterated from 50 to 250 with interval 25 µM. Totally 1000 replications were generated for the global analysis as in **Fig. 5** of the main text. The data files are named as “sample_enzkin_**w**t_noise_data_R_**N**.dat” where N is the replication number taking values from 0 to 999 and w = s or p depending on the trajectory of substrate or product. The first column is time and columns 2 to 10 represents the data corresponding to different initial substrate concentrations s_0_ viz. 50, 75, 100…250 µM. Time course sample datasets on the trajectories of substrate depletion were using in direct nonlinear least square fit with **Eqs. 1** to obtain rate constants [*k*_1_, *k*_−1_, *k*_*2*_] using Marquardt-Levenberg algorithm as depicted in methods section. Using these fit values K_M_ and v_max_ were calculated as *K*_*M*_ = (*k*_−1_ + *k*_*2*_)⁄*k*_1_ and *v*_*max*_ = *k*_*2*_*e*_0_. Analysis over 1000 such time course datasets, one obtains 1000 such values of [*k*_1_, *k*_−1_, *k*_*2*_, *K*_*M*_, *v*_*max*_] which were used to construct the histogram as depicted in **Figs. 6**. Trajectories on the product evolution were used for the steady state analysis by computing average velocities (p / t) at a given reaction time across various substate concentrations and replications.

**Fig. 2E**: “sample_enzkin_**p**t_noise_data_R_**0**.dat”, first and second columns were used as time course data (s_0_ = 50 µM) to NLS fit of (p, t).

**Fig. 2F**: “sample_enzkin_**s**t_noise_data_R_**0**.dat”, first and second columns were used as time course data (s_0_ = 50 µM) to NLS fit of (s, t).

### Sample MATLAB code for NLS fit of time course (p, t) data to Eqs. 1 of main text

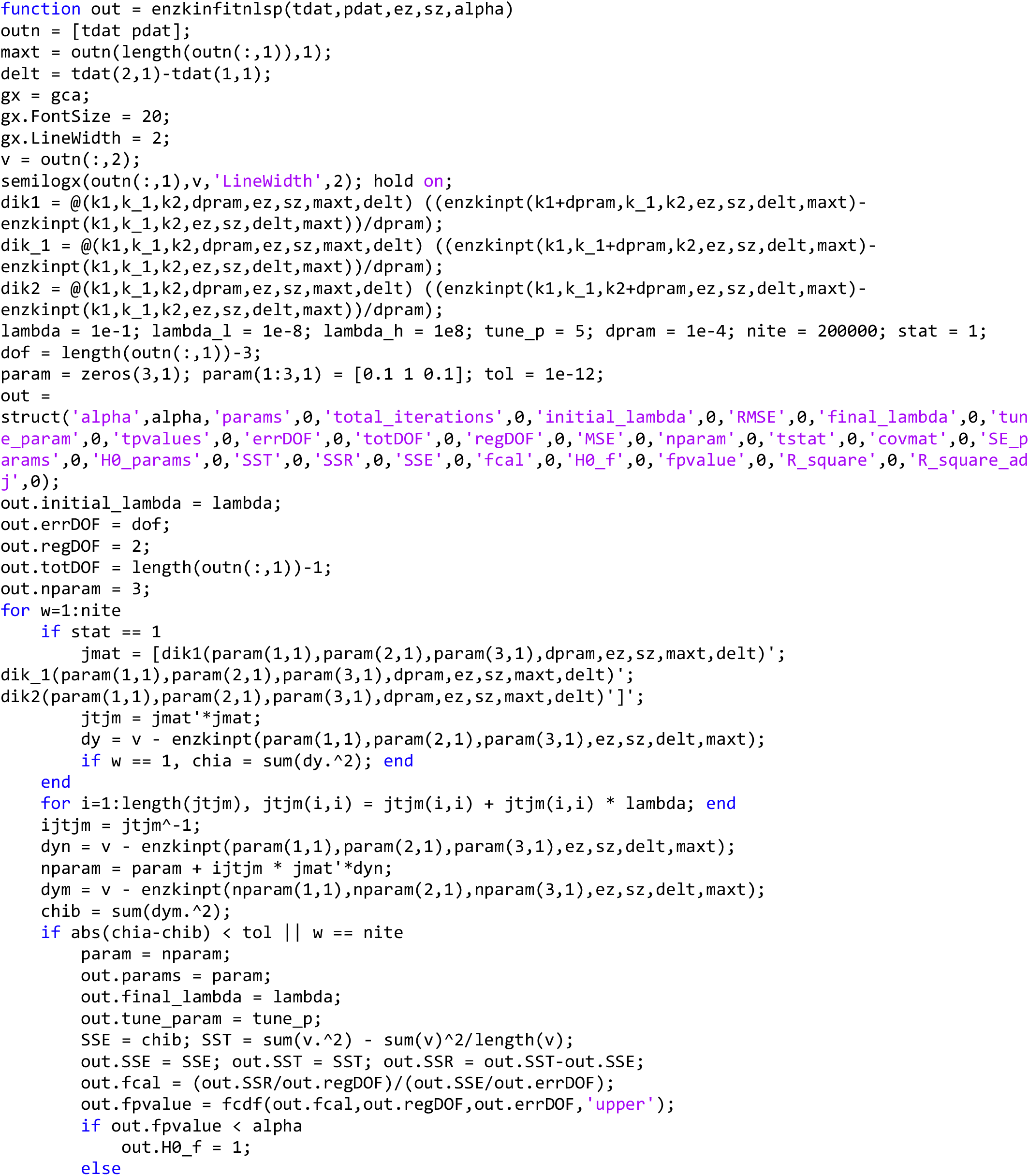
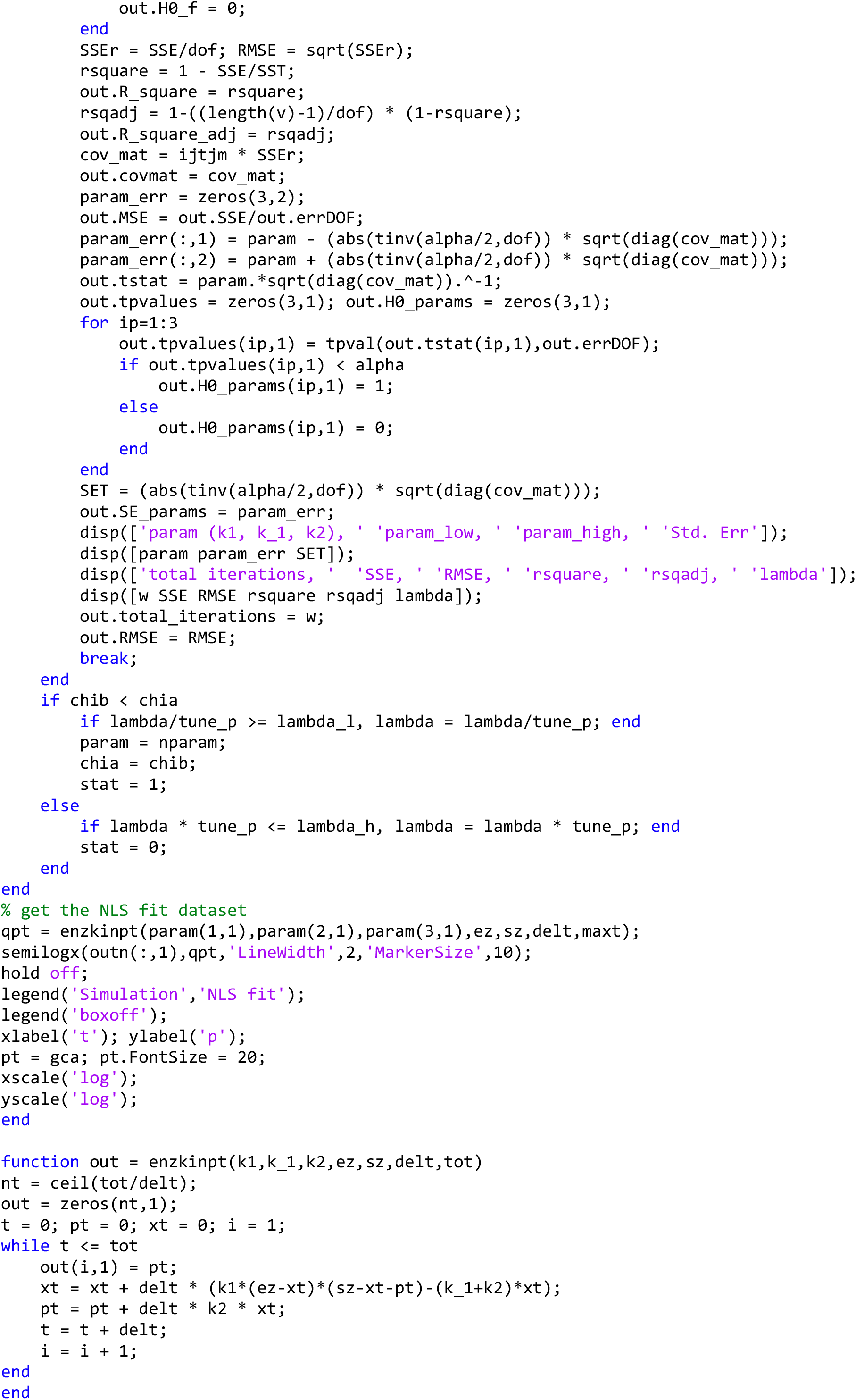

### Sample MATLAB code for NLS fit of time course (s, t) data to Eqs. 1 of main text

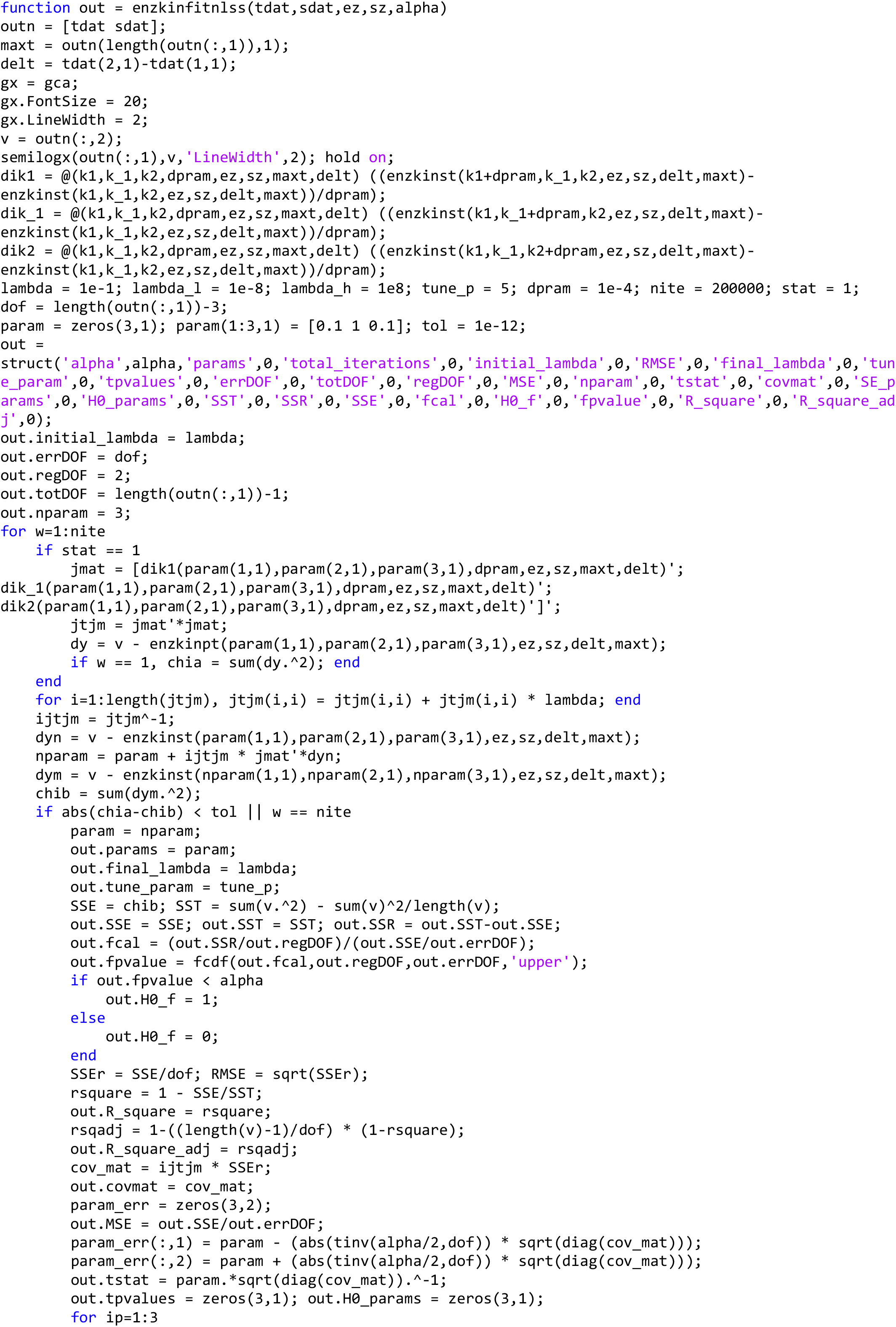
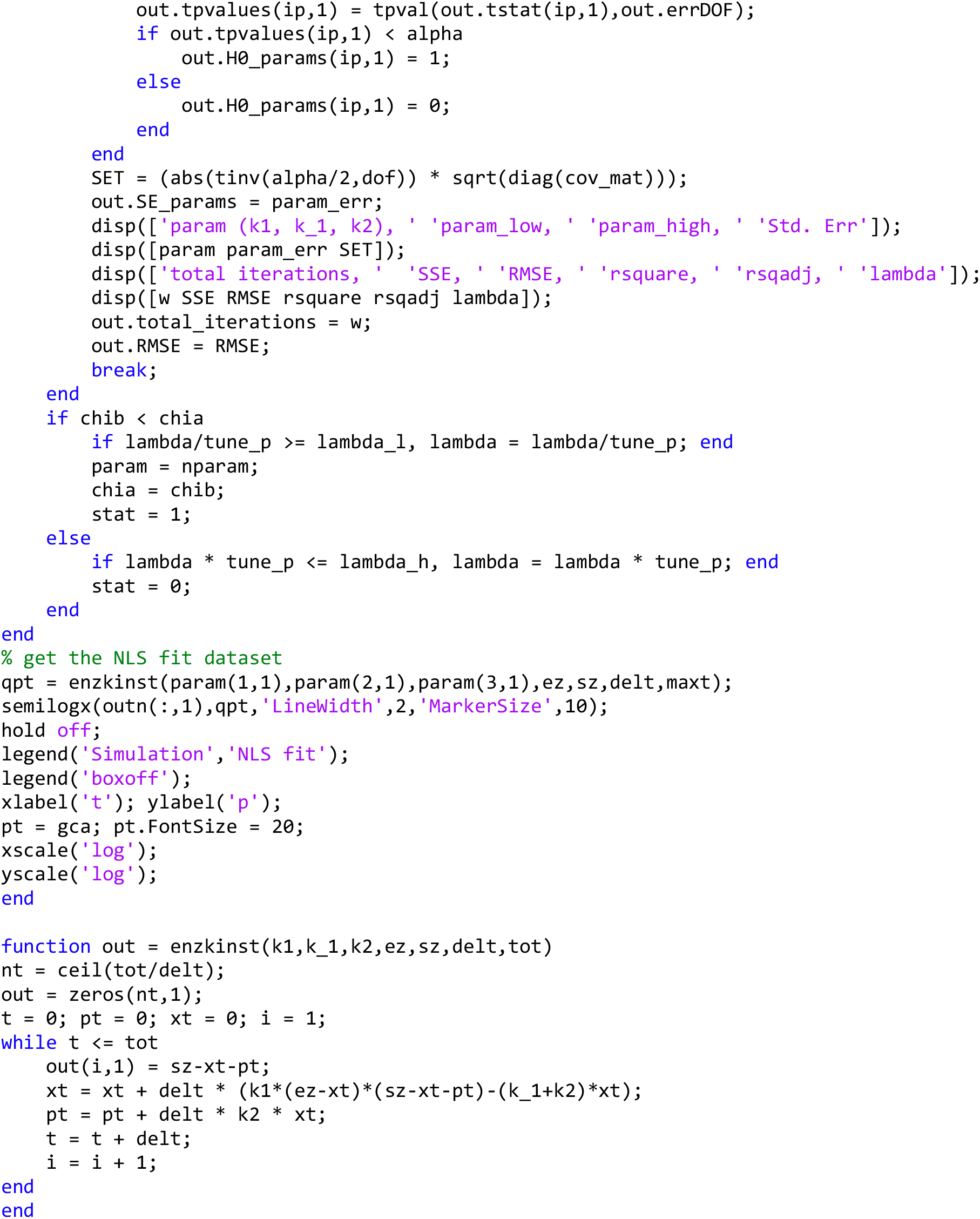

